# Epistatic interaction of the genes *Raw1* and *Raw7* controls barb size and frequency in barley awn roughness

**DOI:** 10.64898/2026.04.27.721039

**Authors:** Muhammad Awais, Matthias Jost, Muhammad Khan, Hélène Pidon, Srijan Jhingan, Axel Himmelbach, Robert Hoffie, Götz Hensel, Jochen Kumlehn, Twan Rutten, Michael Melzer, Jochen C. Reif, Martin Mascher, Nils Stein

## Abstract

Awns of wild barley (*Hordeum vulgare* ssp. *spontaneum* L.) are rough by default due to silicified upward-oriented trichomes on the awn’s epidermis, forming a ratcheted surface, which is advantageous for seed dispersal and burial. Cultivated barley, however, may carry smooth awns covered by smaller barbs or lacking barbs completely. The gene *Raw1* on chromosome 5H is a major factor controlling barley awn roughness and was shown to encode a *LONG AND BARBED AWN1 (LABA1)* homolog. Here we report, by using quantitative analysis of the barb trait, map-based cloning and Cas9-mediated gene knock-out, a second gene *Raw7*, located on barley chromosome 7H, encoding a putative two-component response regulator. We propose that *Raw7* acts downstream of *Raw1* in a cytokinin signaling pathway underlying cell cycle control in epidermal barb primordia cells. *Raw1* and *Raw7* show epistatic interaction, suggesting that *Raw1* acts as the primary driver of barb initiation, while *Raw7* modulates barb size and frequency. Our findings provide the foundation to study the selection and domestication history of the awn roughness trait in barley, and thus to dissect if awn roughness is providing an advantage in cultivated barley or if the trait persisted after domestication due to linkage drag.

**Summary:**

- The presence of silicified upward-oriented trichomes or barbs arising from the epidermis of barley awns is a prominent trait. They form a ratcheted surface which is advantageous for seed dispersal and burial and defense against herbivory.
- Previous work identified *Raw1* on chromosome 5H as a major determinant of awn roughness in barley. Here, we identify a second awn roughness gene, *Raw7* on chromosome 7H, combining quantitative phenotyping of barb traits, map-based cloning and Cas9-mediated targeted mutagenesis for functional analyses.
- Genetic and functional evidence suggests a complex epistatic interaction in which *Raw1* primarily drives barb formation and *Raw7* fine-tunes endoreduplication-dependent epidermal cell expansion and patterning in barb primordia cells. *Raw1* and *Raw7* likely act in a cytokinin-dependent two-component signaling pathway, where *Raw1* promotes local cytokinin activation and *Raw7*, a type-B response regulator, mediates downstream transcriptional responses.
- The proposed pathway suggests additional undetected loci may contribute to awn roughness, and emerging barley pangenome and pan-transcriptome resources provide a framework to identify and functionally validate new candidates.

## Introduction

A key morphological characteristic of barley are its long awns, filamentous distal extensions of the lemma, which are thought to be homologous to a derived or reduced leaf blade as evidenced by mutants like *leafy lemma* (Philipson 1934). As photosynthetically active structures, barley awns play a crucial role in grain development, particularly during the late grain-filling stage (Abebe et al., 2009). Awns in cereals bear upward oriented hook-like single-celled trichomes called barbs, which are primarily distributed along the edges of the awns and characterized by highly silicified cell walls (Hua et al., 2015). The development of barbs is likely driven by endoreduplication-based elongation of epidermal cells (Kondorosi et al., 2000) that are also prevalent on leaves, stems and reproductive organs across various plant families (Peter et al., 1995).

In barley, barbs are observed on the awn surface, causing a rough, ratchet like texture with barb density and size varying significantly among different varieties. Rough-awned barley typically exhibits a higher density of larger barbs along the entire length of the awn, from the base to the apex. In contrast, smooth-awned varieties demonstrate gradual variations in barb size and density or even a complete absence of barbs (Harlan 1920). The rough surface of the barley awns plays a crucial role in seed dispersal over greater distances by attaching to passing animals and adhering to soil (Yuo et al., 2012). This enhances reproductive success by increasing the likelihood of seed germination in new habitats. Additionally, the abrasive texture and sharp edges of rough awns can deter herbivores from foraging, thereby reducing the risk of damage to the crop (Yuo et al., 2012). However, the presence of rough awns in cultivars may pose disadvantages as the abrasive nature of rough awns may contribute to irritation or injury to livestock (Karren et al., 1994) and can cause respiratory issues as a result of extensive inhalation of grain dusts during manual harvest or when using combines (Manfreda et al., 1986). Despite the reported adverse properties, rough-awned barley is globally the predominantly cultivated form (Milner et al., 2019). However, the evolutionary and agronomic significance of barbed awns under current modern agricultural practices remains debated. Some studies suggested a positive correlation between awn roughness and yield, hinting at the historical preference for rough-awned barley varieties (Woodward 1949). In addition, barb formation in barley is reported to positively correlate with the growth of stigmatic hair. The length and density of these filamentous structures are crucial for the success of pollination and seed set, thus a reduction in stigma hair per unit surface area of the ovary could lead to a decreased fertility rate (Harlan and Martini 1940, Zhang et al., 2024, Whitford et al., 2026).

Genetic studies have identified key regulators of barb formation in different plant species. A *LONELY GUY* (*LOG*) family gene named *LONG AND BARBED AWN 1* (*LABA1*) is responsible in rice for the development of long and barbed awns (Hua et al. 2015), as mutations in this gene result in shorter, barbless awns. *LABA1* encodes for a cytokinin (CK) activating enzyme, hence may be involved in the local activation of CK, setting the signaling cascade to cell cycle control, endoreduplication and cell expansion. GWAS demonstrated that awn roughness in barley is under control of at least two major loci on chromosomes 5H and 7H; the gene on 5H, *ROUGH AWN 1* (*Raw1*), was shown to encode for a homolog to rice *LABA1* (Milner et al., 2019). In eggplant (*Solanum melongena*), loss of prickles is linked to mutations in a *LONELY GUY* (*LOG*) family gene, which is involved in CK biosynthesis and organ development (Satterlee et al., 2024). Furthermore, in this study it was demonstrated that *LABA1* homologs or orthologs are involved in the development of prickles in diverse plant species (Satterlee et al., 2024).

The persistence of rough awns in cultivated barley, despite extensive selective breeding, raises two critical questions: why has awn roughness been retained and which genetic mechanisms govern this trait? To address this, our study focused on isolating the gene of the previously reported locus on chromosome 7H (Milner et al., 2019), to further unravel the genetic control and regulatory pathway of barb formation in barley. We utilized high-throughput phenotyping, genetic mapping and validation by Cas9-mediated targeted mutagenesis to uncover the molecular genetic basis of awn barb formation. We identified the second gene regulating awn roughness on barley chromosome 7H (named ‘*Raw7’*), which encodes for a putative two-component response regulator gene, supporting the hypothesis for an interactive role of both genes *Raw1* and *Raw7* as putative components of the same CK signaling pathway. This discovery enhances our understanding of the genetic control of awn roughness in barley as a foundation for studying its evolutionary and ecological significance and as the basis for identifying additional targets in future barley breeding strategies.

## Results

### Contrasting barb size is associated with varying nuclear DNA content in barb cells

To examine whether the contrasting barb phenotypes of the parental genotypes used in our study, ‘cv. Barke’ (2-rowed, rough awned) and ‘cv. Morex’ (6-rowed, semi-smooth awned), are associated with differences in nuclear DNA content, DAPI fluorescence was measured in barb nuclei. DAPI fluorescence intensities revealed higher DNA content of barb cell nuclei in rough-awned Barke (4C) compared to the semi-smooth Morex (2C) (Supplementary Table 1, Supplementary Fig. 1), supposedly the result of endopolyploidization as the functional basis of cell size increase.

### *Raw7* encodes a putative two-component response regulator controlling awn roughness

Previously, a genome-wide association study (GWAS) revealed two major loci associated with awn roughness on chromosomes 5H and 7H (Supplementary Fig. 2) in barley (Milner et al., 2019). One of these segregated in a biparental population of the cross between cultivars Barke x Morex which enabled the positional cloning of the gene *Raw1* on chromosome 5H (Milner et al., 2019). Morex is a ‘semi-smooth’ awned cultivar, as it still develops smaller and less frequent barbs on its awns. Since only one locus segregated in the Barke x Morex cross, we postulated that the second locus on chromosome 7H would segregate in a cross between the semi-smooth cultivar ‘Morex’ and any genotype carrying completely smooth awns. Such genotypes can be found in germplasm collections like the Federal *ex-situ* Genebank of Germany at the Leibniz Institute of Plant Genetics and Crop Plant Research, e.g. the accession MHOR 597 (doi: 10.25642/IPK/GBIS/1470531). Morex and MHOR 597 both carry the homozygous recessive allele of *Raw1* at the chromosome 5H locus as verified by sequence analysis, thus the observed differences in the awn roughness trait are expected to depend on the segregation of an independent genetic factor. In 173 F_2_ plants from the cross Morex x MHOR 597, 19 were scored as smooth-awned and 154 as rough-awned based on haptic assessment. A more objective high-resolution microscopy approach (named ‘optic assessment’) confirmed 53 smooth-awned and 120 rough-awned plants (Fig. 1a). A Pearson’s chi-squared (x*²*) test compared observed F_2_ segregation with Mendelian ratios assuming a monohybrid (*x* locus) or dihybrid (*xy* loci) cross with a dominant-recessive inheritance pattern at 5% significance threshold. Haptic assessment deviated significantly from both the 3:1 monogenic (x² = 18.13) and 15:1 digenic (x² = 6.62) ratios, thereby rejecting the null hypothesis. In contrast, optic observation confirmed a 3:1 monogenic inheritance (x² = 2.93) but differed from the 15:1 digenic ratio (x² = 175.64), clearly indicating the segregation of a single major gene correlated with awn roughness (Table 1).

**Fig. 1.**
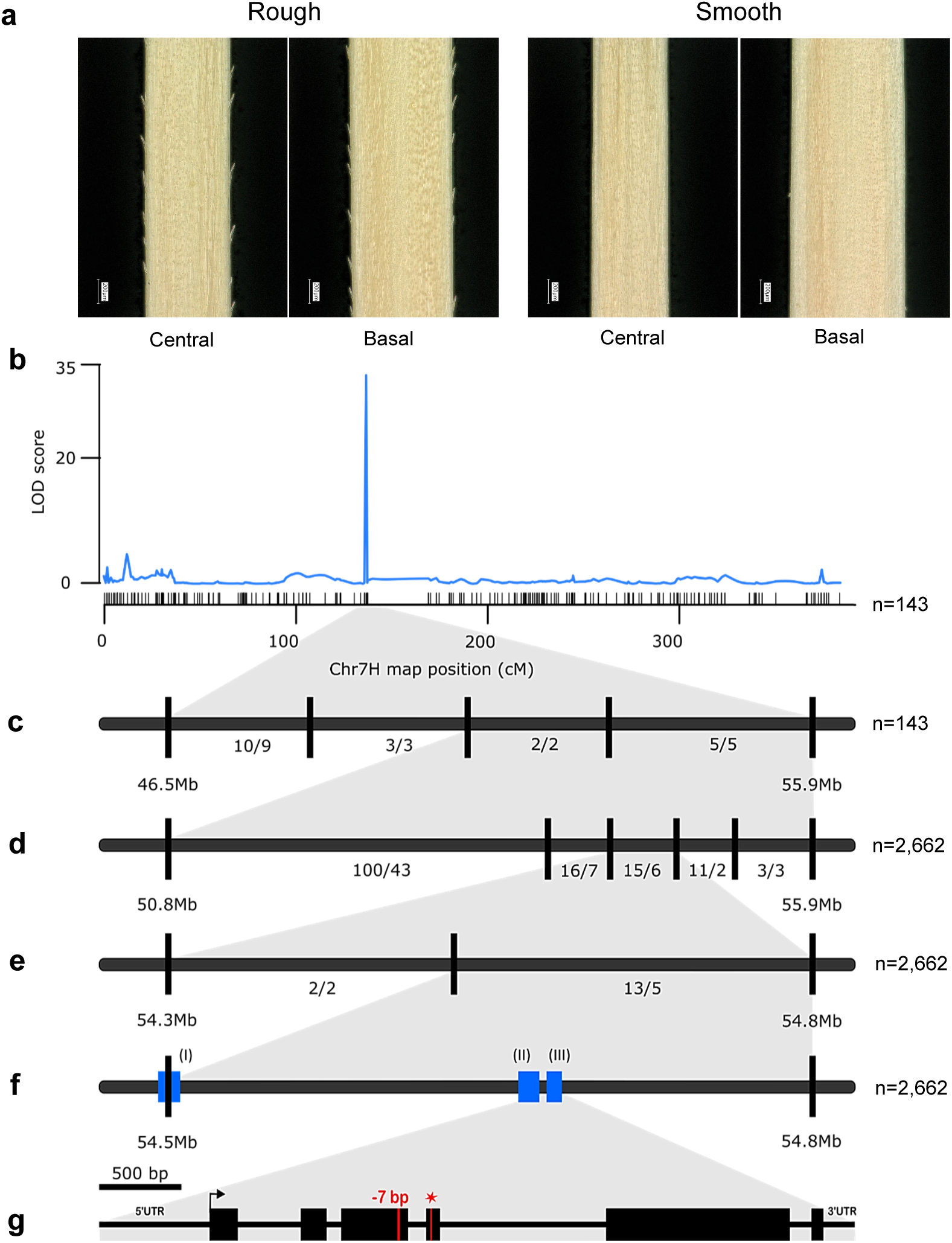
High resolution mapping (HRM) of awn roughness for identification of the gene *Raw7* in the Morex × MHOR 597 F₂ population. **a)** Representative digital microscopic images showing optically assessed rough and smooth-awned F_2_ individuals from the mapping population, at central and basal awn positions. **b)** A single locus associated with awn roughness identified on chromosome 7H using genotyping-by-sequencing. SNP markers are shown as short vertical black bars. **c-e)** Iterative enlarged views (grey shaded areas) of the target locus during HRM with KASP markers (black bars) to identify informative recombinants. The number of phenotyped recombinants is indicated between markers. **f)** HRM defined a 0.3 Mb interval containing three annotated high-confidence genes I, II and III (blue boxes). **g)** Gene model of the strongest candidate (gene III). A seven bp deletion in the smooth awned mutant MHOR 597 (red vertical bar) introduces a premature stop codon (red asterisk) relative to the wild-type cv. Morex. Black boxes indicate exons. Arrow head represents the translation start site. n = number of F_2_ individuals analyzed.

**Table 1.** Assessment of number of genetic loci controlling awn roughness in the Morex x MHOR 597 F_2_ population.

| Awn Phenotype | No. expected |  | Haptic assessment |  |  | Optic assessment |  |  |
| --- | --- | --- | --- | --- | --- | --- | --- | --- |
| | x locus | xy loci | Obs. | $\chi^2$-x | $\chi^2$-xy | Obs. | $\chi^2$-x | $\chi^2$-xy |
| Rough | 129.75 | 162.19 | 154 | 4.53 | 0.41 | 120 | 0.73 | 10.97 |
| Smooth | 43.25 | 10.81 | 19 | 13.60 | 6.21 | 53 | 2.20 | 164.66 |
| Σ (Sum) | 173 | 173 | 173 | 18.13 | 6.62 | 173 | 2.93 | 175.64 |

Observed (Obs.) counts for rough- and smooth-awned phenotypes based on haptic and optic assessments. Expected values and resulting $\chi^2$ statistics are provided for a single (x; 3:1 segregation) and two (xy; 15:1 segregation) loci. Degree of freedom = 1.

For initial low-resolution genetic mapping, we performed genotyping-by-sequencing in 178 F_2_ plants (Pop-1_Batch-1) of the same population. Quality-checked reads were aligned to the MorexV2 reference genome assembly (Monat et al., 2019) with a mapping rate of 97.6%. Variant calling yielded 468,621 raw variants (including 22,342 insertions and deletions). A genetic linkage map was constructed using 1,167 high-confidence single-nucleotide polymorphisms (SNPs) (Supplementary Fig. 3).

Genome-wide QTL analyses using inclusive-composite interval mapping (ICIM) with optically assessed phenotypes identified a single major locus on chromosome 7H exceeding the LOD threshold of 4.23. This locus (LOD = 61.11, *P* < 0.05), designated ‘*raw7HS’*, explained 76.9% of phenotypic variance and was localized to the short arm of barley chromosome 7H (Supplementary Fig. 4). This coincided with the GWAS results of Milner et al. (2019), representing a 4.1 cM genetic and a 9.2 Mbp physical interval (Fig. 1b). For high-resolution genetic mapping (HRM), the interval was equidistantly saturated with KASP markers and the same F_2_ population was screened as well as 16 individuals of each of the 20 recombinant F_3_ families, that had been identified to represent the initial genetic interval (Fig. 1c).

This was followed by iterative cycles of KASP marker design and mapping in 143 F_2_ plants and their respective F_3_ families, previously identified in a reduced 5.1 Mbp physical interval after increasing the genetic resolution by screening 2,662 F_2_ individuals (Pop-1_Batch-2-9) with the eight KASP markers (Fig. d-f). Phenotypes of 61 recombinant F_2_ plants were validated in their respective F_3_ families, which narrowed down the genetic target interval to 0.3 Mbp (Fig. 1e), containing three high-confidence genes (Fig. 1f). Due to their tight genetic linkage with the awn roughness trait and based on the available structural and functional gene annotations, all three genes were qualifying as putative candidate genes (Monat et al., 2019, Mascher et al., 2021). Candidate genes I (*HORVU.MOREX.r2.7HG0546700/HORVU.MOREX.r3.7HG0658780*) and III (*HORVU.MOREX.r2.7HG0546720/HORVU.MOREX.r3.7HG0658800*) were functionally annotated as putative ‘Two-Component Response Regulator’ genes, while the gene II (*HORVU.MOREX.r2.7HG0546710/HORVU.MOREX.r3.7HG0658790*) was classified as a putative ‘Histidine Kinase’. Since the expression of the rice gene *LONG AND BARBED AWN 1* (*LABA1*) (Hua et al., 2015), a homolog to the barley gene *Raw1* (Milner et al., 2019), was correlated with higher expression of a *RESPONSE REGULATOR (RR6)* gene in the rice awn primordia of the wild-type long and barbed awn genotype, we prioritized the candidate genes I and III for further functional analyses. Comparative sequence analysis of the two genes identified a seven base pair deletion in the coding region of gene III of the mutant MHOR 597 compared to wild-type Morex, causing a frame shift and resulting in a pre-mature stop codon (Fig. 1g). Candidate genes I and II did not exhibit any variations at the polypeptide level between wild-type and mutant genotypes. Consequently, *HORVU.MOREX.r3.7HG0658800* was considered as the most plausible candidate gene, acting in the regulatory pathway of awn roughness in barley. To maintain the consistency with established barley gene naming nomenclature (Franckowiak and Lundqvist 2004) we designated the gene formally as *ROUGH AWN7* (*Raw7)*.

### Functional validation of *Raw1* and *Raw7* by Cas9-mediated site-directed mutagenesis

To functionally validate the candidate gene *Raw7* and to determine the single and combined effects of both genes *Raw1* and *Raw7* in an isogenic background, a Cas9-endonuclease based gene editing approach was employed. The *Raw1* target vector ‘pGH469’, carrying a single gRNA cassette, and the *Raw7* target vector ‘pRH48’, carrying four gRNAs, were used for co-delivery of respective agrobacterial transfer DNAs (T-DNAs) to immature embryo explants of cv. Golden Promise [(rough awned: *Raw1*/*Raw1* (*AA*), *Raw7*/*Raw7* (*BB*)] to generate *raw1/raw1/raw7/raw7* (*aabb*) double mutants (experiment prefixed ‘BRH42’). Additionally, we targeted *raw1* knockout lines (BG741_E08 and BG741_E22; Golden Promise background) to edit the *Raw7* locus to independently obtain *aabb* double mutants (experiment prefixed ‘BRH41’). A total of 189 plants (109 for BRH41 and 80 for BRH42) were regenerated (T_0_/M_1_ generation), out of which 173 were confirmed by PCR to carry the T-DNA. In the population derived from the *raw1* mutant background, 82 plants carried mutations in *Raw7*. In the population derived from the wild-type background, 34 plants carried mutations in *Raw7*, 42 in *Raw1*, and 11 plants were mutated in both genes. The regenerated M_1_ plants were analyzed for the presence of induced mutations on one or both gene loci by amplifying both target genes and subsequent Sanger sequence analyses. Selected M_1_ plants were self-pollinated to obtain T-DNA-free segregants with all possible allelic combinations of the two mutated awn roughness genes (*AABB*, *AAbb*, *aaBB* and *aabb*) in the next generation (T_1_/M_2_). The subsequent sequence analysis of both target genes validated the presence of homozygous induced mutations in M_2_ plants (Fig. 2a and Supplementary Table 2).

**Fig. 2.**
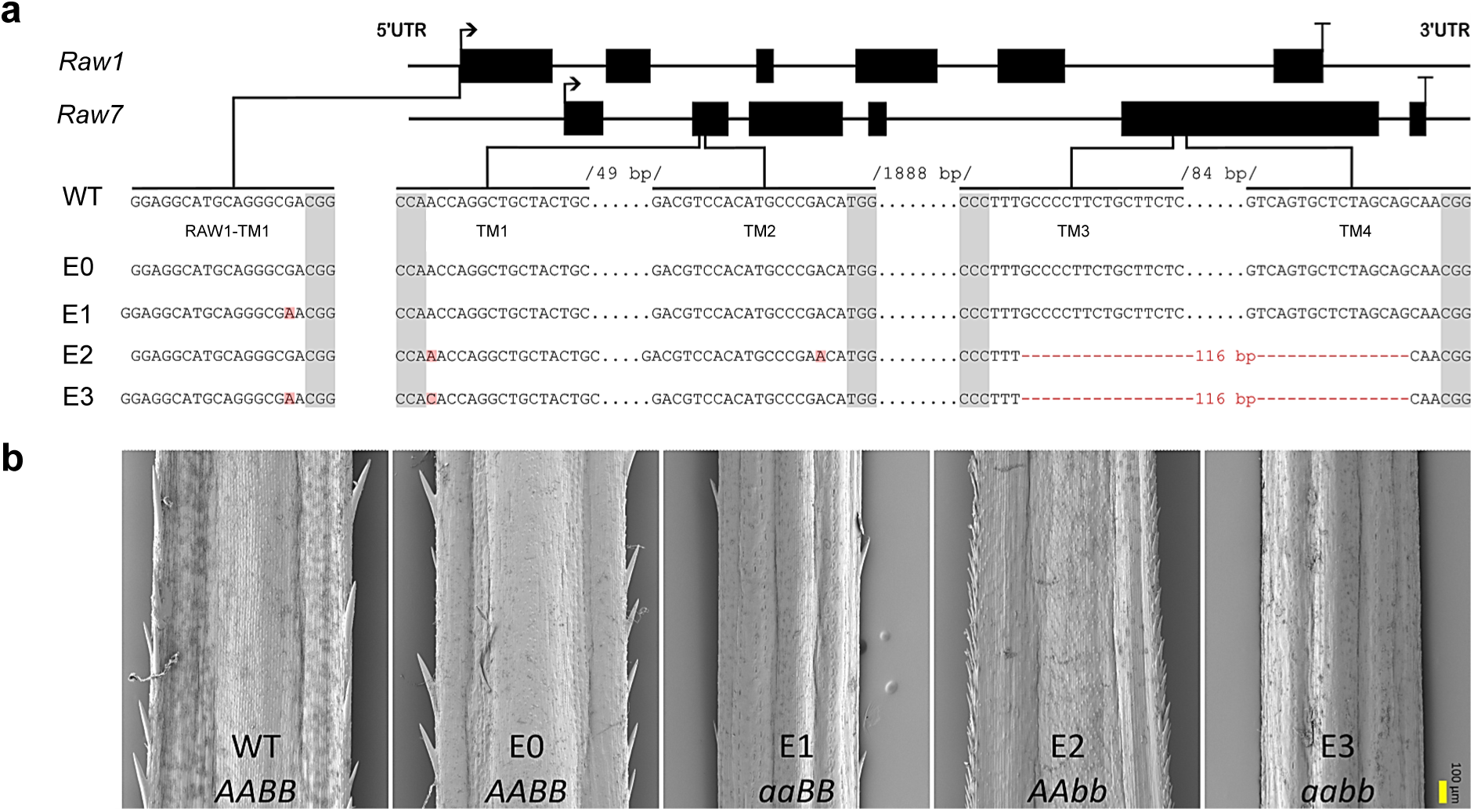
Phenotypic and genotypic characterization of Cas9-induced homozygous mutants of the *Raw1* and *Raw7* genes. **a)** Gene models of both rough awn genes with indicated target sites for the guide RNAs (gRNAs). One (‘Raw1-TM1’) and four (‘TM1-4’) target motifs (TM) were selected for *Raw1* and *Raw7*, respectively (Supplementary Table 9). Target sequence alignments of the wild-type (WT) and the M_2_ plants originating from independent editing events (E0-E3) are shown. Grey shaded bases correspond to the protospacer-adjacent motifs (PAM). Cas9-induced mutations are shown in red. **b)** Awn phenotypes of representative M_2_ plants in homozygous genetic combinations of both targeted genes (*A* = *Raw1*, *a* = *raw1*, B *= Raw7*, *b* = *raw7*). Scale bar (yellow) = 100 µm.

Awn roughness phenotyping was performed for the selected M_2_ progenies. To validate the phenotypic differences in the mutated events (E0-E3), the non-mutated (E0) and a non-transformed/non-regenerant wild-type Golden Promise (WT) were included in the analysis as wild-type controls (Fig. 2a). The phenotypes and genotypes were validated in at least three M_2_ plants per genotype which were confirmed to be T-DNA free (Fig. 2b). The plant category E0, homozygous for wild-type alleles at both loci (*AABB*), exhibited the rough awn phenotype, consistent with the Golden Promise wild-type control (WT). The least rough awn phenotype, comparable to the semi-smooth awns of Morex, was obtained in category E1 (*aaBB*). E2 (*AAbb*) was characterized by small-sized barbs present at a much higher density along the awn surface. The category E3 (*aabb*), homozygous double mutants, displayed a pronounced smooth awn phenotype, showing the complete loss of barbs from the central and basal awn areas. These results demonstrate that *Raw7* is involved in barb formation of barley awns. We provide plausible evidence for the independent and interactive roles of the genes *Raw1* and *Raw7* regulating barb size and density, and thus collectively controlling the quality of awn roughness in barley.

### Awn roughness in barley is controlled by complex epistatic interactions between *Raw1* and *Raw7*

The genes *Raw1* (*AA*) and *Raw7* (*BB*) were shown to contribute distinct effects to barb development and awn roughness in barley. To elucidate the genetic interaction between both genes, comprehensive genotypic and phenotypic analyses were performed using diagnostic KASP marker assays on an F_2_ population (Pop-2) of Barke (*AABB*) x MHOR 597 (*aabb*), segregating for alleles at both loci. A set of 162 F_2_ individuals was identified from the expected nine genotypic classes: 14 F_2_ plants of genotype *AABB*, 26 of each *AaBB* and *AABb*, 20 of *AaBb*, 16 of *aaBB*, 11 of *aaBb*, 12 of *AAbb*, 24 of *Aabb* and 13 of *aabb*. Microscopy-based optical assessment revealed four distinct awn phenotypes across all genotypic combinations. These included the three known parental phenotypes i.e., Barke and Golden Promise as ‘rough-awned’, Morex as ‘semi-smooth-awned’, MHOR 597 as ‘smooth-awned’, and a fourth category displaying a high frequency and density of small-sized barbs, classified as ‘super-rough-awned’ (Fig. 3 and Supplementary Fig. 5). The genotypic classes were phenotypically comparable to those observed in the combined functional validation through site-directed mutagenesis of *Raw1* and *Raw7* (Fig. 2b).

**Fig. 3.**
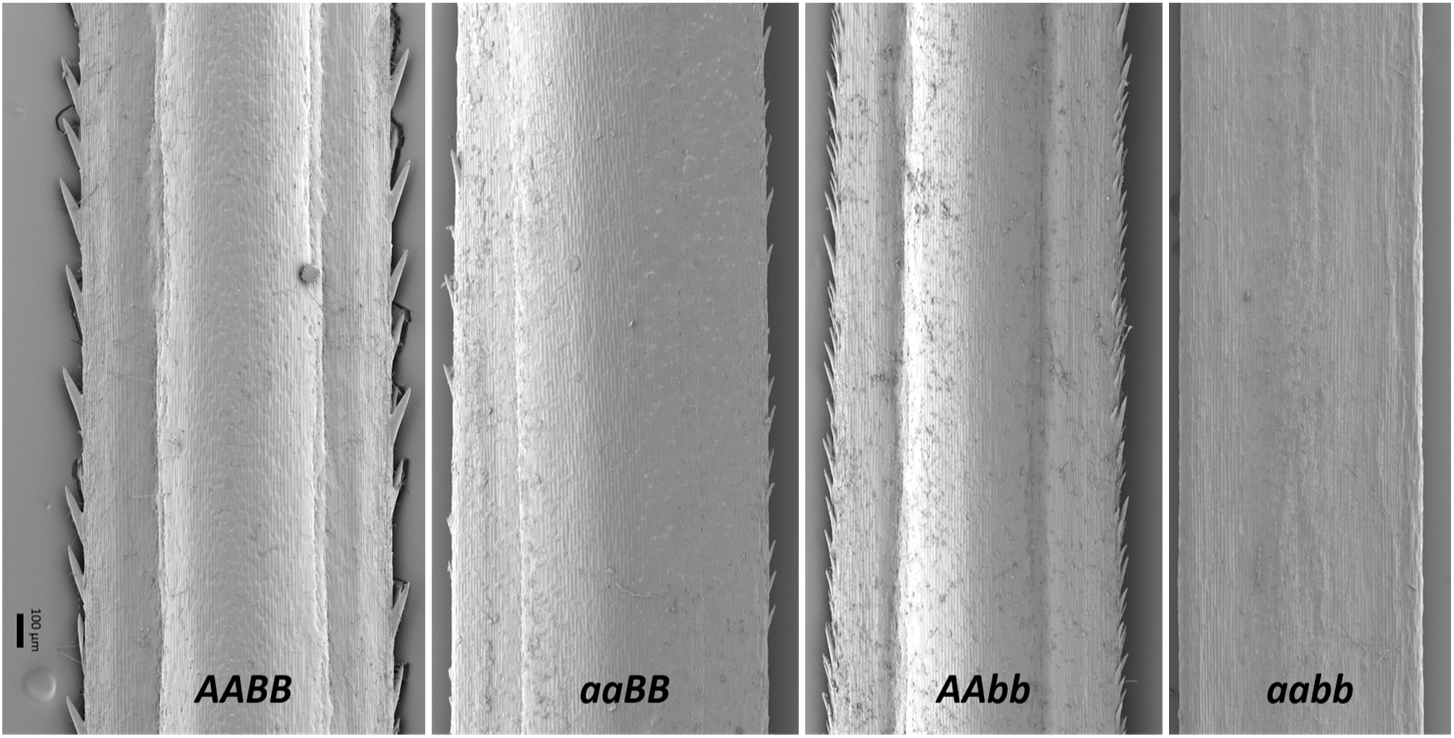
Phenotypic variation in awn roughness among homozygous genotypic combinations of *Raw1* and *Raw7* in Barke x MHOR 597 F_2_ population. Scanning electron microscopic images at the basal awn position depicting four observed phenotypic classes, each corresponding to homozygous allelic combinations of wild-type or mutated alleles for the rough awn genes (*A* = *Raw1*, *a* = *raw1*, *B* = *Raw7*, *b* = *raw7*).

Awn roughness was assessed via manual barb counting and automated image analysis using BarbNet (Narisetti et al., 2023). Although both methods were strongly correlated (r *=* 0.96*, p <* 0.001), the automated image analysis for awn roughness was chosen for subsequent high-throughput analysis (Supplementary Fig. 6). Barb density and size varied significantly (F = 52.28*, p* < 0.001) across *Raw1/Raw7* genotypic classes at basal awn positions (Fig. 4). The highest densities were observed in genotypes carrying at least one wild-type *Raw1* allele in a homozygous *raw7* background (*AAbb* and *Aabb*), while the *aabb* double mutants completely lacked barbs. Genotypes heterozygous at *Raw7* within a homozygous *raw1* background (*aaBb*) did not differ significantly from *aabb*, and all remaining genotypes exhibited intermediate barb density (Fig. 4a). A similar pattern was observed at the central awn position, with significant variation across genotypes (F = 44.7, *p* < 0.001). The highest barb density was again observed in *AAbb* plants, whereas *aabb* plants lacked barbs entirely. The other genotypes showed intermediate barb sizes, consistent with the trends observed in the basal position.

**Fig. 4.**
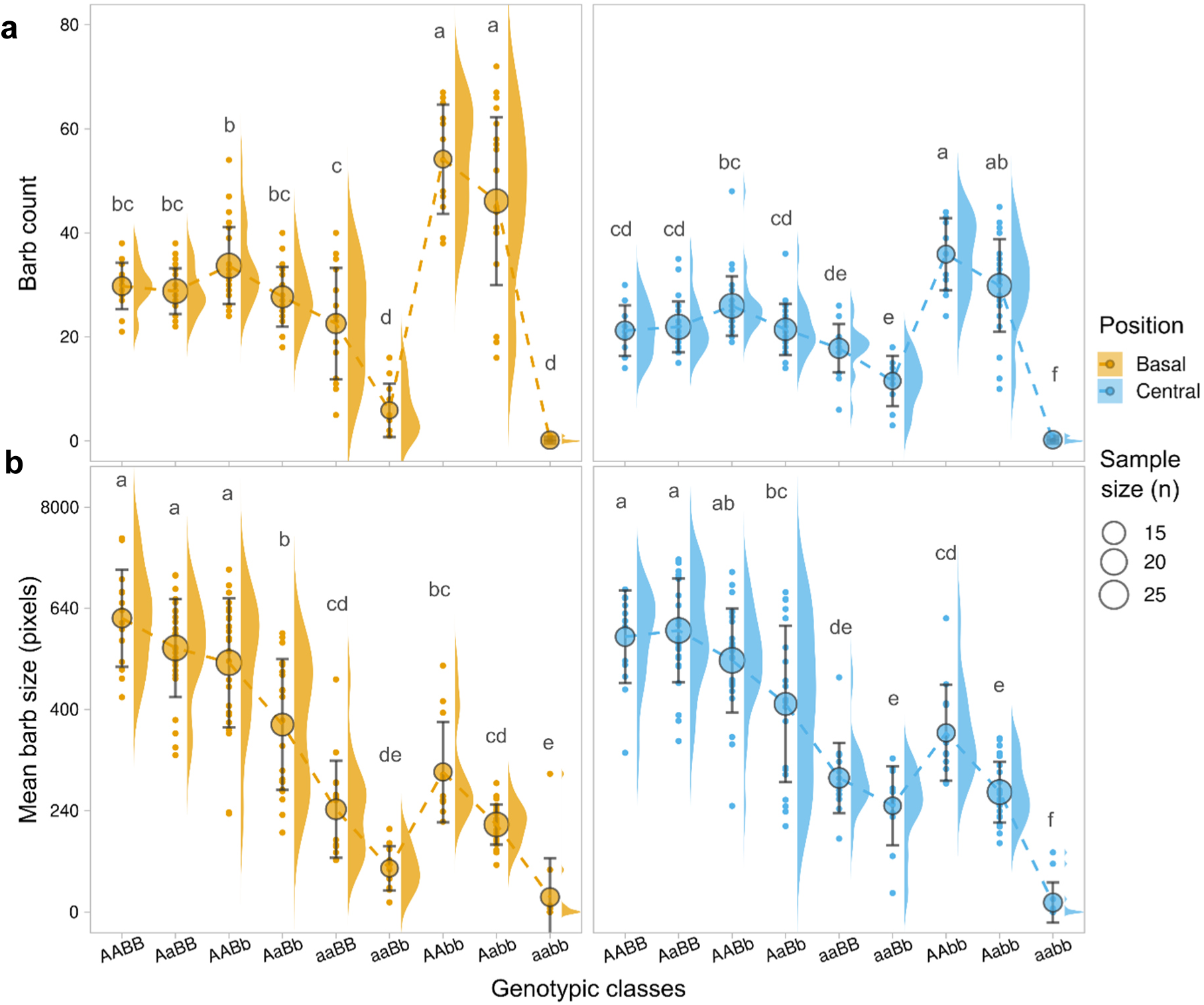
Impact of *Raw1* and *Raw7* in regulating the barb density and size. Automated image analyses via BarbNet showing **a)** barb count and **b)** size at central and basal awn positions across nine genotypes composing wild-type and/or mutated alleles of *Raw1* and *Raw7*. Mean barb size was measured as number of pixels occupied with barbs. Colored dots represent individual data points for respective classes; basal and central awn positions. Larger circles indicate mean values, with sizes proportionate to the sampling size in each genotypic class (*A* = *Raw1*, *a* = *raw1*, *B* = *Raw7*, *b* = *raw7*). Error bars represent standard deviation. The letters above each class indicate statistical groups assigned based on ANOVA (*P* < 0.001) and Tukey’s HSD (*P* < 0.05) analyses.

Barb size also varied significantly among genotypes at both positions. At the basal awn position, ANOVA showed a strong genotype effect (F = 69.2, *p* < 0.001). The largest barbs were observed in genotypes carrying at least one wild-type allele of both genes (*AABB*, *AaBB* and *AABb*), while the smallest barbs occurred in genotypes *aabb* and *aaBb*. The remaining genotypes exhibited intermediate barb sizes. At the central awn position, significant differences were also detected (F = 57.3, *p* < 0.001). The largest barbs occurred in genotypes with at least one wild-type allele of both *Raw1* and *Raw7*, Minor barb measurements in *aabb* likely reflect false-positive detection by BarbNet, rather than true barb formation. Genotypes carrying either gene in homozygous recessive state showed intermediate barb sizes (Fig. 4b).

The roles of *Raw1* (*A*) and *Raw7* (*B*) were tested with ANOVA and regression models that partitioned additive (Ad) and dominance (Dom) effects and their interactions. At the basal awn position, both genes and most interactions significantly affected barb count (Supplementary Table 3). The exceptions were interactions involving *Raw7* dominance (Ad-A x Dom-B and Dom-A x Dom-B), which were not significant, in line with the shared statistical grouping of *aaBb* with *aabb* phenotypes. Basal barb size was similarly influenced by both genes individually and in combination, except for Dom-B (F = 2.66, *P* = 0.11) and the Dom-A x Dom-B interaction (F = 0.03, *P* = 0.86), which did not contribute significantly (Supplementary Table 4). At the central awn position, all genotypes significantly affected barb number, where *Raw1* showed strong additive and dominance effects (*P* < 0.001) and *Raw7* showed moderate but significant effects (*P* < 0.05) (Supplementary Table 5) with the Ad-A x Dom-B interaction being weaker (F = 2.76, *P* < 0.1). Central barb size was influenced by both genes independently, with limited interaction effects except for Dom-A x Ad-B (F = 7.47, *P* < 0.01), which had a moderate effect (Supplementary Table 6). Taken together, our results suggest *Raw1* as a major determinant of barb density, whereas *Raw7* modulates it such that a single wild-type *Raw7* allele reduces barb density. For barb size, both genes act complementarily when wild-type alleles at both loci produce the largest barbs, while loss-of-function at either locus reduces size. These patterns were consistent at both basal and central awn positions, demonstrating complex independent and interactive roles of *Raw1* and *Raw7* in controlling awn roughness in barley.

## Discussion

### *Raw1* and *Raw7* are factors of a putative two-component signaling system in barb primordia

This study aimed to identify the genetic factor underlying a second major locus on barley chromosome 7H, contributing to the control of awn roughness, in addition to the gene *Raw1,* both previously identified through GWAS (Milner et al., 2019). *Raw1*, located on chromosome 5H, was identified encoding a putative cytokinin riboside 5ʹ-monophosphate phosphoribohydrolase (Milner et al., 2019). Here we demonstrate that the second gene *Raw7,* allocated to chromosome 7H, encodes for a putative two-component response regulator (RR), an ortholog to gene *OsRR22*, a subfamily-I type-B RR with nuclear localization in rice (Li et al., 2019). *LABA1*-regulated RRs have been shown to play a role in barbed awn development (Hua et al., 2015). These RRs are components of two-component signaling systems, which help cells to respond to environmental stimuli through phosphorylation-mediated signal transduction involving histidine kinases and RRs (Ito and Kurata 2006, Tsai et al., 2012). While the functional annotation of *Raw7* implies the involvement in such a pathway, further studies are required to validate its precise biological function.

### Proposed regulatory model for barb formation in barley

Homology-based functional annotation suggests that *Raw1* and *Raw7* are likely to act in the same signaling pathway, regulating barb formation in barley awns. Barbs are single-celled trichomes originating from epidermal cells that are significantly larger than typical epidermal cells. We demonstrated through densitometric measurements of nuclear DNA content that barb precursor cells in barley awn primordia underwent endopolyploidization, a modified cell cycle in which DNA replication occurs without mitosis or cytokinesis, which promotes cell differentiation and expansion (Takahashi and Umeda 2014, Fambrini and Pugliesi 2019, Pinto et al., 2024). Specialized cells like trichomes, root hairs, and fruit tissues commonly undergo this process, forming polyploid nuclei to induce and promote cell growth (Liu et al., 2016). Trichome development typically occurs in three stages: initiation, endoreduplication, and expansion, regulated by genetic and hormonal factors (Fambrini and Pugliesi 2019). Phytohormones play a key role in regulating these phases. For instance, cytokinin (CK) stimulates trichome formation by regulating genes such as *ZFP6*, *ZFP8*, and *GIS2* (Gan et al., 2007, Zhou et al., 2013). Gibberellins (GA), CK, and jasmonic acid (JA) regulate trichome morphogenesis through genes like *GL1*, *GL3*, *TTG1*, and *TRY* (Pesch et al., 2014), whereas salicylic acid (SA) inhibits trichome initiation (Traw and Bergelson 2003). Auxins influence glandular trichome differentiation in tomatoes by interacting with CK and GA to regulate development and ensure proper trichome spacing (Payne et al., 2000). CK plays a central role in trichome and barb development, often interacting with auxins (Tang et al., 2023). In rice, functional studies of *LABA1* or *An-2*, another LOG family gene, have confirmed their enzymatic role in CK biosynthesis, converting inactive CK nucleotides into bioactive CKs such as isopentenyladenine (iP) and trans-zeatin (tZ), ensuring a local pool of bioactive CK for downstream signaling (Kurakawa et al., 2007, Gu et al., 2015, Hua et al., 2015). In Arabidopsis, CK modulates the transition from mitosis to endoreduplication through the transcription factor Arabidopsis Response Regulator 2 (ARR2), which activates the expression of CCS52A1, a key regulator of the endocycle (Larson-Rabin et al., 2009, Takahashi and Umeda 2014, Fathy et al., 2022).

Based on existing models of trichome development involving CK signaling, a similar mechanism may regulate barb formation in barley. Our findings on the nature and functional annotation of *Raw1* and *Raw7* align with the broader CK signaling pathway described in Arabidopsis (Kieber and Schaller 2018), where bioactive CKs are perceived by histidine kinase (HK) receptors, and a two-component signaling system activates CK-responsive Response Regulators (RRs) in the nucleus. Extrapolating this model to barley (Fig. 5) puts the gene *Raw1* at the beginning of the signaling cascade, leading to localized active CK levels, which eventually trigger *Raw7* to regulate downstream genes involved in cell cycle control. The consequence is localized cell-specific endoreduplication and cell expansion, ultimately leading to barb formation in barley awns.

**Fig. 5.**
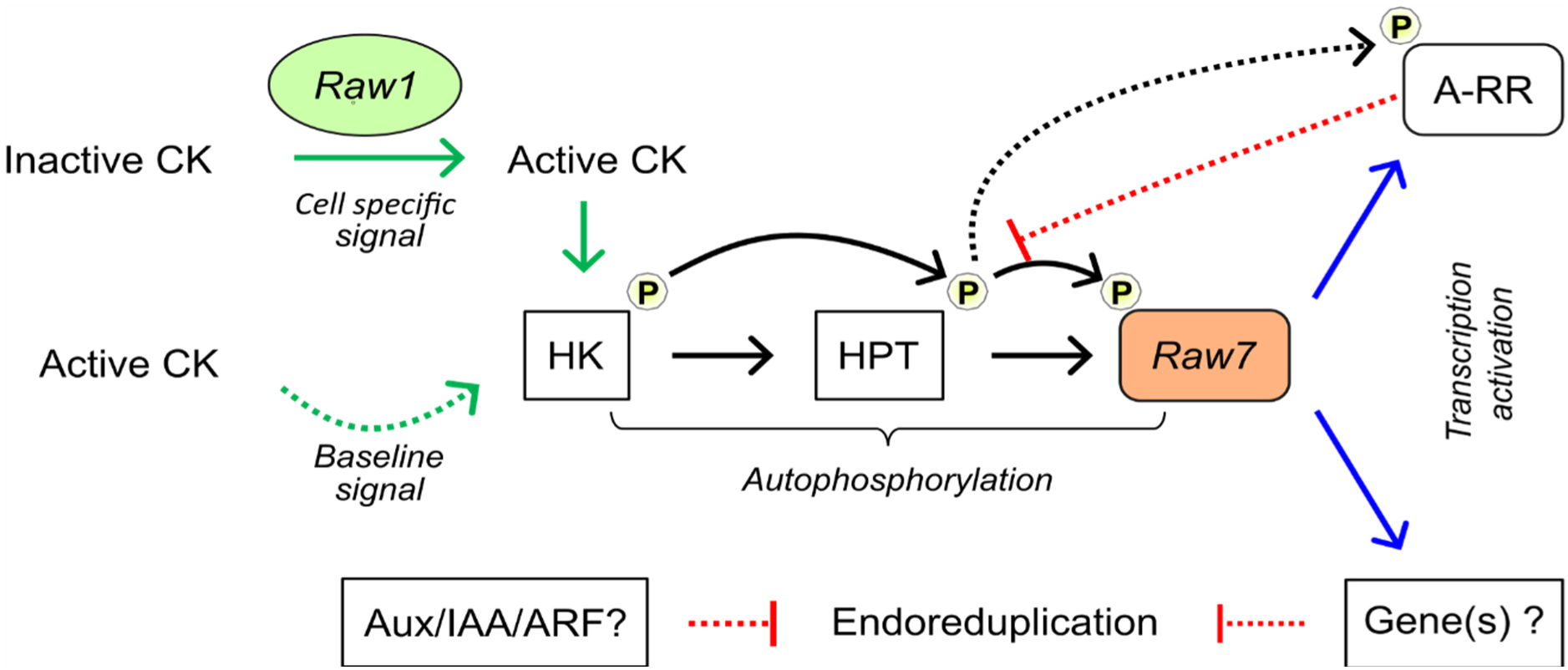
Proposed signaling model for regulation of awn roughness in barley. Based on Arabidopsis models, a cell-specific cytokinin (CK) signal is initiated by *Raw1* (LOG family). This signal is perceived by Histidine Kinase (HK) and transferred via Histidine Phosphotransfer Protein (HPT) to activate Raw7 (Type-B Response Regulator) through phosphorylation. Once active, Raw7 regulates the transcription of downstream factors, including a Type-A Response Regulator (A-RR) that provides negative feedback. Raw7 may also interact with auxin-responsive genes (Aux/IAA/ARF) to modulate the cell division machinery, triggering endoreduplication and subsequent barb extension.

If the control of barley awn roughness follows such a gene regulatory pathway model, it implies that additional factors are likely involved and could thus be discovered. The earlier GWAS analysis for awn roughness had revealed only two major loci controlling awn roughness in barley (Milner et al., 2019). Additional loci may have escaped the detection limit due to low allele frequencies in the studied barley population. However, functional annotation of the barley pangenome (Jayakodi et al., 2024), as well as the availability of extensive pan-transcriptome data (Guo et al., 2025) might provide new opportunities to systematically survey for functional sequence diversity in predicted genes of the postulated pathway. This may allow prioritizing candidates for Cas9-mediated site-directed mutagenesis and functional verification in the future.

### Genetic interaction between *Raw1* and *Raw7*

Our proposed model for regulation of barley awn roughness suggests a coordinated interaction of *Raw1* and *Raw7* where the distinct phenotypes represent specific physiological consequences of modified cell cycle and patterning. While *Raw1* initiates the process, the complex epistatic interactions observed with *Raw7* suggests it may act as a regulator for barb density likely by restricting the differentiation of epidermal cells into barbs and enforcing spacing between them. This may be accompanied by the successful transition from standard mitosis to the endoreduplication as a means for localized DNA replication needed for cell expansion as observed in our densitometric data (To and Kieber 2008, De Veylder et al., 2011, Breuer et al., 2014). The loss of functional alleles at either locus likely disrupts the CK mediated signaling cascade, preventing the nucleus from reaching the high endopolyploid thresholds necessary to drive the formation of large barbs. The “smooth” or “semi-smooth” phenotypes in mutant lines represent a failure to complete this modified cell cycle, whereas the “super-rough” phenotype represents an uncontrolled initiation phase followed by restricted expansion and disrupted patterning. This may reflect a loss of neighbor inhibition, leading to reduced spacing between barbs and a decoupling of initiation and expansion processes (Pesch and Hülskamp 2009).

### Future directions for understanding the ecological and agronomic dynamics of awn roughness in barley

The presence of awns in barley, whether rough or smooth, carries significant ecological and agronomic importance (Abebe et al., 2009). Given the prevalence of rough-awned modern barley cultivars, the role of roughness in barley awn morphogenesis raises a critical question: do rough-awned barley varieties hold advantages over smooth-awned ones, despite their adverse properties during crop harvest? The advantages of rough awns, akin to trichomes in other plants, are well-documented. DeWitt et al., (2023) demonstrated that rough awns in wheat improved ecological fitness by aiding in cooling under heat stress. The role of trichomes in defense mechanisms against insect herbivores by creating physical barriers that impede movement, feeding, or oviposition and facilitation of seed burial and protection against predation and environmental hazards like grassland fires (Wood and Lenné 2018) have been linked as essential for survival and eventual domestication. However, domesticated barley faces challenges distinct from its wild ancestors (Yuo et al., 2012). In large-scale agriculture, rough awns can complicate grain handling and storage, increasing the risk of spoilage and contamination (Harlan 1920). Additionally, in livestock farming, rough awns compromise animal health by causing irritation or injury when used as feed (Karren et al., 1994).

The identification of *Raw1* and *Raw7* as important regulators of awn roughness presents an opportunity to further explore these agronomic trade-offs. Marker-assisted selection could enable the development of smooth-awned cultivars without compromising fertility or yield since associations between rough awns and fertility, seed set, and ultimately yield have been reported (Zhang et al., 2024). To contextualize these findings within barley evolution, selective sweep analysis for uncovering domestication patterns across diverse genotypes could ascertain if the rough awned trait resulted from artificial selection, genetic drift or as an artifact of linkage drag with other agronomically relevant genetic factors. Furthermore, a putative pleiotropic effect that phenotypically links rough awns with stigma hairiness could serve as a proxy for yield stability (Woodward 1949, Whitford et al., 2026). Ultimately, comprehensive studies on the link between awn roughness on yield given modern agricultural practices could clarify whether the rough awned phenotypes offer a competitive edge, bridging the gap between ancestral ecological benefits and the precision required for modern barley optimization.

## Materials and methods

### Plant material and growth conditions

The semi-smooth-awned cv. Morex was crossed with the smooth-awned mutant MHOR 597 - doi: 10.25642/IPK/GBIS/1470531 (Scholz and Lehmann 1958) to generate an F_2_ population designated as ‘Pop-1’. The population was divided into two batches for low- and high-resolution mapping named ‘Pop-1_Batch-1’ and ‘Pop-1_Batch-2-9’ with 192 and 2,662 individuals, respectively. A second F_2_ population derived from the cross between the rough-awned cv. Barke and the smooth-awned mutant MHOR 597 was named ‘Pop-2’ representing 368 F_2_ plants. Seeds were sown in 96-welled pots and leaf samples were collected at two weeks after sowing for DNA isolation. Seedlings were transplanted into pots (16 cm for Pop-1_Batch-1, and Pop-2 and 9 cm or 48-well plates for Pop-1_Batch_2–9) for awn phenotyping and grain harvesting. Recombinant plants identified in low-resolution mapping were propagated to the F_3_ generation to generate recombinant inbred lines. All plants were grown under greenhouse conditions (18 °C day/14 °C night, 16 h light). The wild-type rough-awned cv. Barke, semi-smooth-awned cv. Morex and the smooth-awned mutant MHOR 597 were grown as parental controls in all experimental batches. After growth *in vitro*, all regenerants from Cas9-mediated targeted mutagenesis were transferred to greenhouse conditions (20 °C day/18 °C night, 16 h light). Plants were propagated to the M_2_ generation for the selection of particular genotypes as well as for phenotyping and grain harvesting.

### Genomic DNA (gDNA) extraction and quantification

For gDNA extraction, six-centimeter leaf segments were collected from two to three weeks old seedlings. A GTC-NaCl (Guanidinium thiocyanate - Sodium chloride) protocol was followed (Milner et al., 2019). DNA quality was visually inspected using 1% (w/v) agarose gel electrophoresis with ethidium bromide staining (Sharp et al., 1973). Gel standards included lambda DNA (20, 50, 100 ng) and a 1 kb O’GeneRuler (Thermo Scientific, USA). All gDNA samples were diluted 1:200 with de-ionized distilled water before quantification. DNA concentration was estimated using a NanoDrop spectrophotometer (Thermo Scientific, USA) and a Qubit 2.0 Fluorometer (Invitrogen, USA) with the Broad Range assay kit. Qubit working solutions and standards (0 and 100 ng/μL) were prepared as per the manufacturer’s guidelines.

### Phenotypic analysis of awn roughness

Phenotyping for awn roughness was performed on apical, central and basal segments of awns from mature spikes. For qualitative analyses, a haptic assessment was performed by sliding the fingers from the awn apex to the base. For awns with haptic resistance, the “rough” category was assigned, whereas awns with no resistance were considered as “smooth”. For quantitative measurements of barb counts, awn images (2.32 mm and 3.48 mm of awn per position) were acquired using a digital microscope (KEYENCE DEUTSCHLAND GmbH, Germany) at 100x or 150x magnification with consistent resolution settings. In a high-throughput quantitative assay, digital microscopy-based images were processed with the automated image analysis tool BarbNet (Narisetti et al., 2023). The mean values from three awns per primary spike were taken for each plant. Images were quality controlled for focus, resolution and contrast. Images that deterred reliable image analyses for barb count, density and size quantifications were removed.

For scanning electron microscopy analysis of awn phenotype, freshly isolated awns were fixed for 72 h at 8 °C in 4% formaldehyde in 50 mM phosphate buffer pH 7.0. After dehydration in a graded ethanol series, samples were critical point dried in a Bal-Tec CPD 030 critical point dryer (Bal-Tec AG, Switzerland). Following, probes were placed onto carbon adhesive discs, gold coated in an Edwards S150B sputter coater (Edwards High Vacuum Inc., UK) and examined in a Zeiss Gemini300 scanning electron microscope (Carl Zeiss Microscopy GmbH, Germany) using 5 kV acceleration voltage. Images were stored as .TIFF files.

### Genetic mapping of *Raw7*

For low-resolution mapping (LRM) of the awn roughness trait, the mapping population Pop-1_Batch-1 was genotyped via Genotyping-by-Sequencing (GBS) using the protocol of Wendler et al. 2014. Pooled libraries were size-fractionated (150-600 bp) using a Blue Pippin™ instrument (Sage Science, USA) and assessed on the Agilent TapeStation 4200 (Agilent Technologies, USA) before sequencing on the Illumina HiSeq2500 platform (Illumina Inc., USA). Raw reads in the .fastq format were trimmed using Cutadapt v1.15 (Martin 2011) to remove adapters and aligned to the MorexV2 reference genome (Monat et al. 2019) using BWA-MEM v0.7.17 (Li and Durbin 2009). Resulting files were generated and sorted using Novosort (https://www.novocraft.com/documentation/novosort-2/). Variant calling was performed with SAMtools v1.7 (Danecek et al., 2021) and BCFtools v1.6 (Danecek et al., 2021), generating a raw variant calling format (.vcf) file. To yield high-confidence single nucleotide polymorphism (SNP) markers, the raw .vcf file was filtered for biallelic SNPs with mapping quality ≥40, read depth ≥5 reads per position, missingness at thresholds 10% and 20%, and minor allele frequency ≥5%.

High-confidence SNP markers from the GBS dataset were used to construct a genetic linkage map via JoinMap v4.1 (Van Ooijen 2006) employing the Kosambi mapping function to estimate distances in centiMorgans (cM). SNPs were grouped into seven linkage groups, one for each of the seven barley chromosomes based on recombination frequencies (<0.45) and logarithm of odds (LOD) scores (>1). Monomorphic markers and those with segregation distortions were excluded to ensure map accuracy. Double crossovers were manually corrected to avoid artefacts caused by sequencing errors.

Quantitative trait locus (QTL) analysis for awn roughness was conducted using inclusive composite interval mapping (ICIM) in QTL ICiMapping v4.1.0.0 (Li et al., 2008, Meng et al., 2015) with a walking speed of 1 cM. Using high-confidence SNP markers from genetic linkage mapping and the phenotypic data were as inputs, marker-trait associations were identified using 1,000 permutation tests to determine the LOD threshold. Mapping was performed using the Haley-Knott regression on the “R/qtl” package (Broman et al., 2003) on R within R Studio (R Core Team 2020).

### SNP genotyping using Kompetitive Allele-Specific PCR (KASP) assays

High-resolution mapping (HRM) of the rough awn locus and genotyping of Pop-2 were performed using KASP assays (Supplementary Table 7 and Supplementary Table 8). Flanking SNP markers identified through linkage mapping were converted into KASP assays (3CR Biosciences, UK). Sequence data of SNPs, with 50 bp up- and downstream, were extracted using the Integrated Genome Viewer (IGV) v2.16.1 and used to design allele-specific primers. KASP reactions were set up in 384-well plates, indexed, and run on the ABI 7900HT RT-PCR system (Thermo Scientific, USA), following the manufacturer’s guidelines (PACE Quick reference Guide v1.4). Fluorescent signals were analyzed using SDS v2.4.1 software (Thermo Scientific, USA) to determine allele-specific amplifications.

### Targeted mutagenesis of genes controlling awn roughness trait

To validate genes controlling awn roughness in barley, Cas9-mediated mutagenesis was performed using the CasCADE modular vector system as described by (Hoffie 2022). Binary vectors pGH469 and pRH48 were used to knockout *Raw1* and *Raw7*, respectively.

For *Raw1*, the pGH469 binary vector contains a single RAW1-TM1 gRNA cassette (GenBank: HG793095.1) run by the *OsU3* promoter, which was cloned by *Bsa*I into pSH121 (Gerasimova et al., 2018).The resulting Cas9 (under control of the *ZmUbi1* promoter) and gRNA expression cassettes were cloned into the binary vector backbone p6i-2×35S-TE9 (DNA Cloning Service, Hamburg, Germany) based on *Sfi*I sites. The resulting pGH469 was used for *Agrobacterium*-mediated barley transformation as described by (Marthe et al., 2014). For this, cultivar ‘Golden Promise’ was used due to its wild-type alleles at both loci and its high transformation and regeneration efficiency. Transgene-free, homozygous M_2_ lines were selected, namely BG741-E08 and BG741-E22, carrying independent mutations events of 1 bp insertions.

For *Raw7*, gRNAs were designed using the whole-genome sequence of ‘Golden Promise’ (Schreiber et al., 2020) according to Kumlehn et al., (2018). The online tool CRISPRDB (https://crisprdb.org/) was used to screen for putative target motifs in exons 2 and 5, ensuring specificity and proper SpCas9 PAM sequence (5’-NGG-3’). Two pairs of target motifs were selected based on criteria previously described (Koeppel et al., 2019) including the secondary structure of the cognate gRNAs as predicted by RNAfold (http://rna.tbi.univie.ac.at/) and the distance between the paired target motifs. Off-target analysis was performed with BLASTn on Geneious Prime v2023.1.2 (Biomatters Ltd., New Zealand). The intermediate vector pMA7 was cloned according to the CasCADE protocol with four gRNA expression cassettes under control of the *TaU6* promoter and *Cas9* under control of the *ZmUbi1* promoter. All together were cloned into the binary vector backbone p6i-2×35S-TE9 (DNA Cloning Service, Hamburg, Germany) based on *Sfi*I sites. The pRH48 vector was introduced into ‘Golden Promise’ wild-type in co-transformation with pGH469 (BRH42) as well as to BG741-E08 and -E22 *raw1* mutant lines (BRH41) via *Agrobacterium*-mediated transformation, and T-DNA-positive plants were selected during regeneration using hygromycin. These primary mutants/transgenics (M_1_/T_0_) were genotyped by PCR for the presence of the T-DNA and PCR amplicons of the target regions of *raw1* and *raw7* were Sanger sequenced.

To identify transgene-free, homozygous mutants, gDNA was extracted from 20-40 progeny per selected primary mutant plant and tested for the presence of the T-DNA using PCR with specific primers for the *hpt* and *cas9* transgenes (Supplementary Table 9). PCR was performed to amplify the transgene fragments using GoTaq^®^ G2 DNA polymerase (Promega GmbH, [M7845]) and QuantiTect^®^ SYBR^®^ Green PCR kit (QIAGEN, 204141) following the manufacturer’s guidelines in a PCR machine (Alpha cycler 4, PCRmax™ or TGradient, Biometra GmbH). PCR products were then visualized on 2% (w/v) agarose gel to verify presence/absence of T-DNA fragments. Editing patterns were confirmed by PCR-amplifying the target regions (Supplementary Table 10). Amplicons were purified using NucleoFast 96 PCR plates (Macherey-Nagel GmbH &Co. KG, Germany) and the sample concentration was measured using a Nanodrop device (Thermo Scientific, USA) following the manufacturer’s instructions. The purified PCR products were sent for Sanger sequencing (Supplementary Table 11), performed by a service provider (LGC Genomics, Germany) following their guidelines. The resulting sequence files were aligned with the wild-type Golden Promise using the alignment tool on Geneious Prime v2023.1.2 (Biomatters Ltd., New Zealand) to confirm mutations in selected T-DNA-free plants.

### Statistical analyses of phenotypic data

Pearson’s chi-squared (x*²*) test was used to compare observed F_2_ segregation with Mendelian ratios assuming a monohybrid (*x* locus) or dihybrid (*xy* loci) cross with a dominant-recessive inheritance pattern (*p* < 0.05; degree of freedom = 1).

Barb quantification data obtained from BarbNet were further analyzed in RStudio (R Core Team 2020). Analysis of Variance (ANOVA) and post-hoc Tukey-HSD tests were applied based on pairwise significant differences in awn roughness among the nine genotypic classes *AABB*, *AaBB*, *AABb*, *AaBb*, *aaBB*, *aaBb*, *AAbb*, *Aabb* and *aabb*). To assess epistatic interactions, the following linear regression model was used:

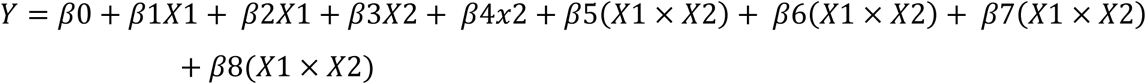

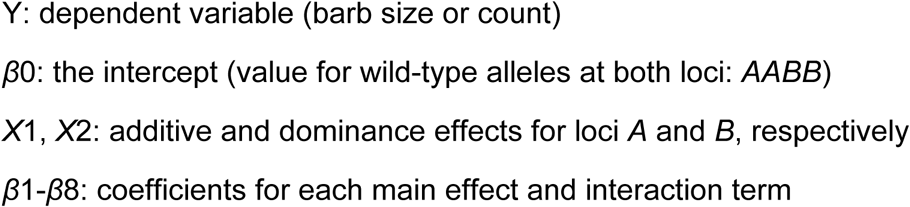

## Acknowledgments

We thank Mary Ziems and Dagmar Svoboda for assisting in all greenhouse and growth chamber experiments for plant and seed material production. We also acknowledge Jacqueline Pohl and Susanne König for providing excellent technical support during DNA sequencing experiments and Sibille Freist for the generation of the *raw1* mutant plants.

## Author contributions

NS and MJ conceived the study. NS supervised the study together with HP. MA, MJ and MK performed experiments and MA analyzed the data. SJ and MA jointly wrote the manuscript. GH designed and cloned the *raw1*-related constructs and genotyped the M_0_ and M_1_ progeny for this target. JK conceived and supervised the generation of genome-edited plants and RH performed the transformation experiments. TR and MM performed all microscope-based analyses. Sequencing data were produced by the IPK NGS-team under the guidance of AH. JCR guided the epistasis analyses. M Mascher provided support during bioinformatic analyses. All authors revised and approve the manuscript.

## Data availability

The Genotyping-by-Sequencing dataset from the Morex x MHOR 597 F_2_ population for low-resolution mapping is available in the European Nucleotide Archive (ENA), accessible under the identifier PRJEB102976. Scripts used for data analyses are available at https://github.com/oves1/barley-awn-roughness-genetics.

---

**Supplementary Fig. 1.**
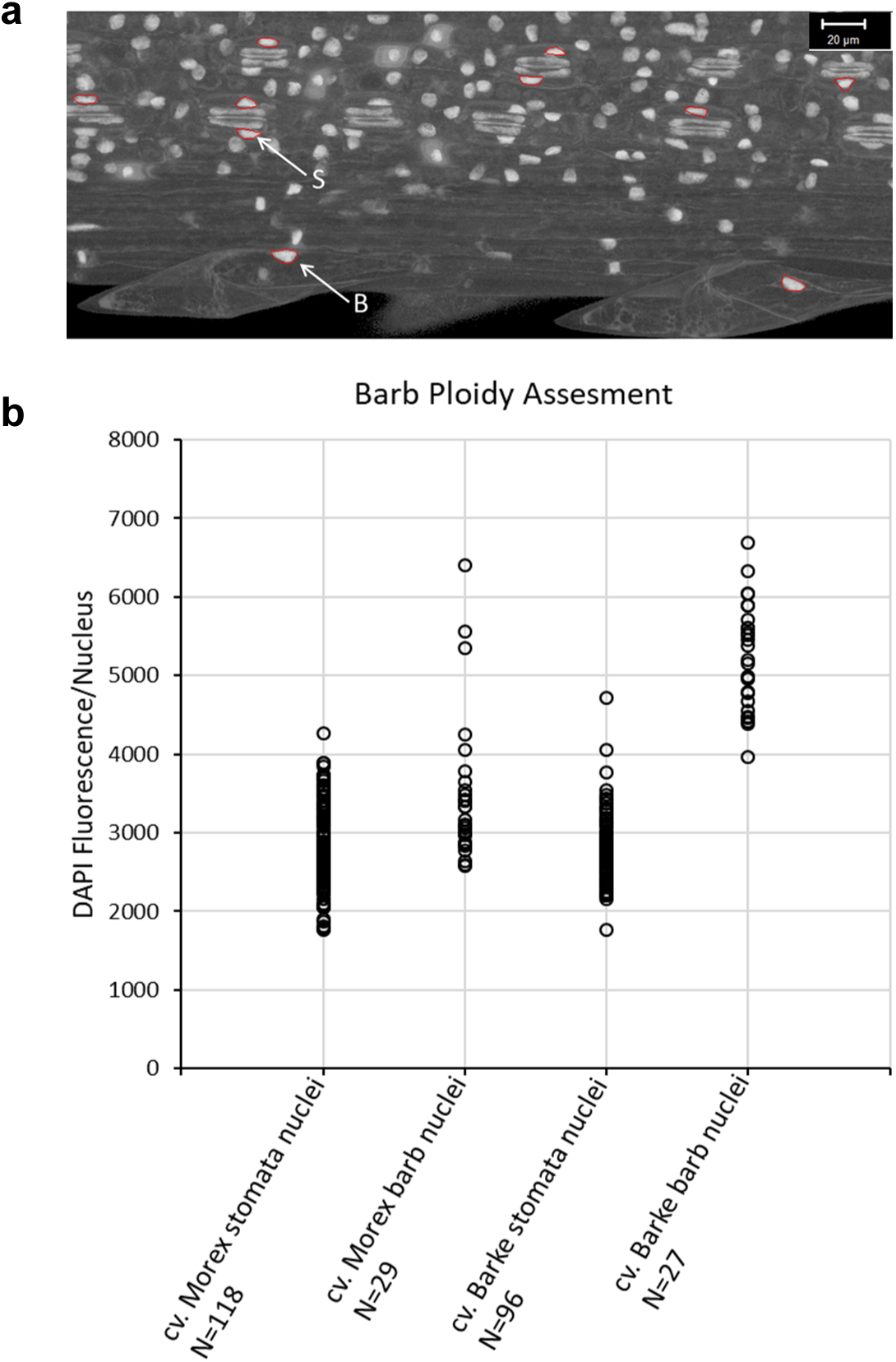
Ploidy assesment of barb nuclei in the awns of cv. Morex and cv. Barke after DAPI staining. **a)** Confocal image showing detail of awn from cv. Barke stage W8.0 with brightness adjusted to distinguish stomata guard cell nuclei (S) and barb (B) nuclei. **b)** Measurements of absolute DAPI fluorescence intensity. Stomata guard cell nuclei used as an internal control for 2C ploidy. N = number of nuclei analyzed.

**Supplementary Fig. 2.**
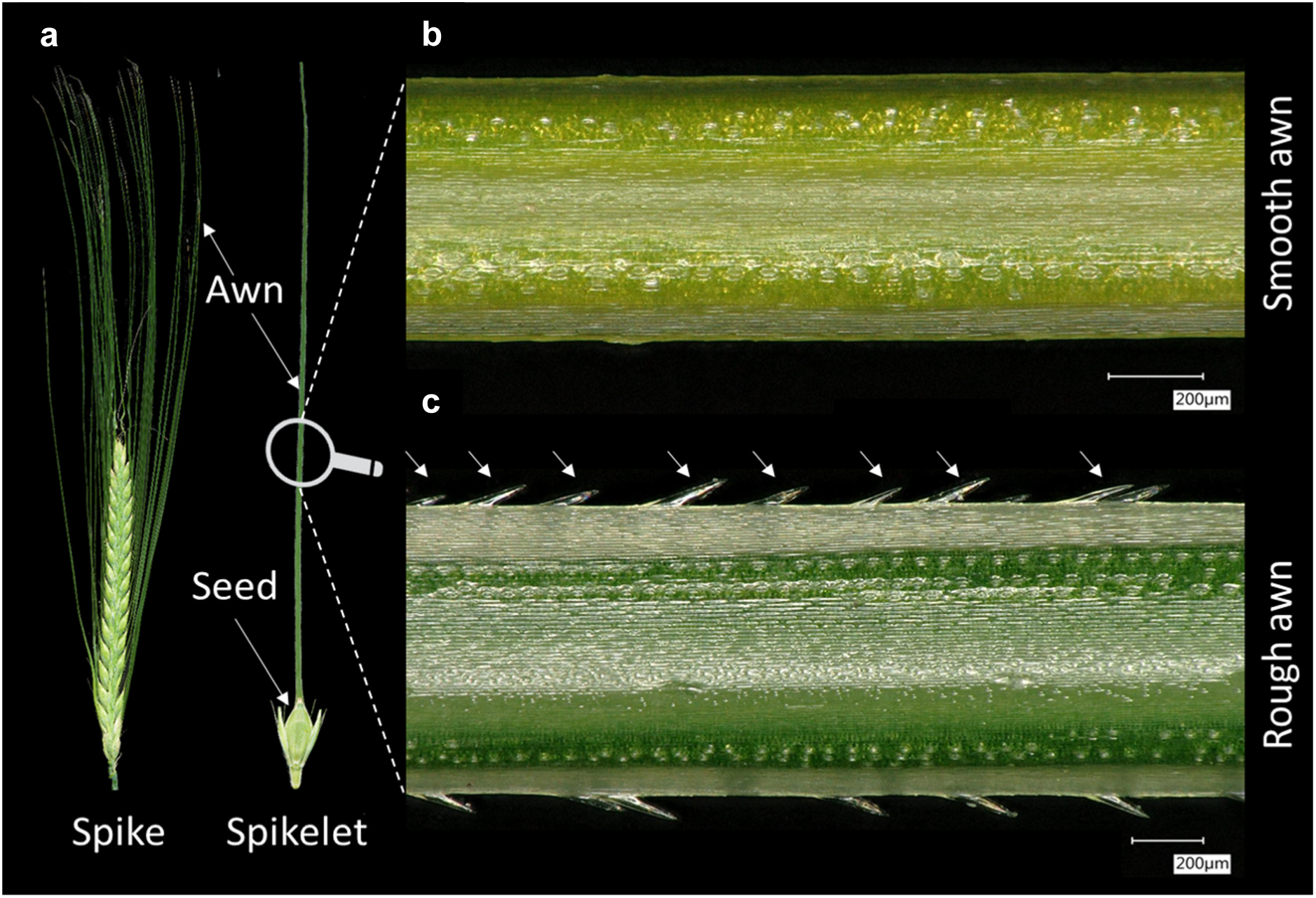
Representative images showing barley spike and awn morphology. **a)** The two-rowed type barley spike and its spikelet bearing awns. Digital microscopy image (150x magnification) showing central parts of two barley genotypes **b)** without (smooth) or **c)** with (rough) barbs (marked with white arrows).

**Supplementary Fig. 3.**
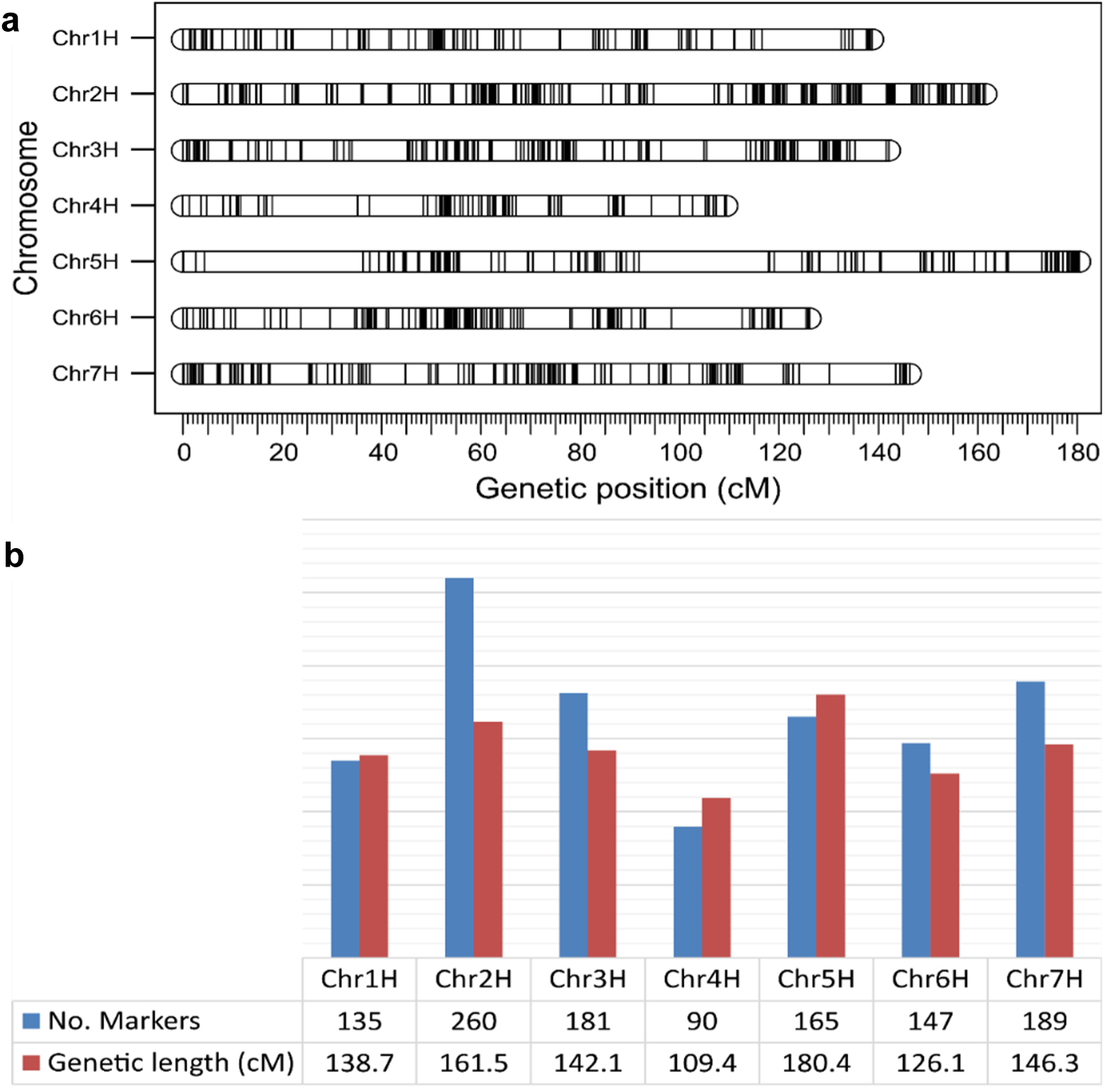
Genetic mapping using genotyping-by-sequencing (GBS) for the F_2_ population originating from the Morex x MHOR 597 cross. **a)** The genetic linkage map shows the positions of GBS identified SNP markers (black vertical bars) along seven barley chromosomes (Chr1H-7H). The marker coordinates are shown in centiMorgans (cM). **b)** Summary statistics of the genetic linkage map constructed using GBS identified high-confidence SNP markers.

**Supplementary Fig. 4.**
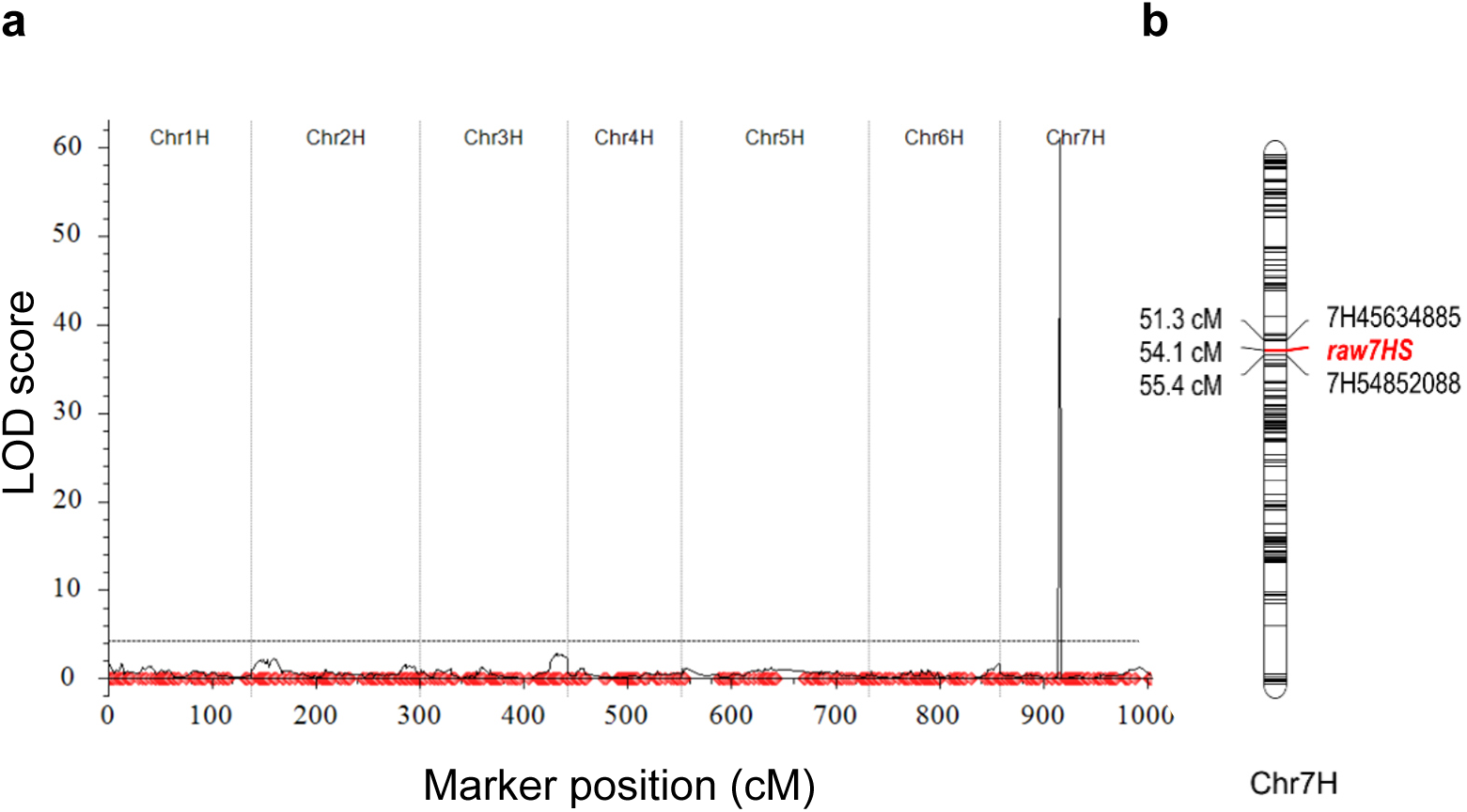
QTL analyses scans from the Morex x MHOR 597 F_2_ mapping population (Pop-1_Batch-1) for **a)** the whole genome showing a single significant (LOD threshold in black dotted horizontal line at 4.23) peak on chromosome 7H and **b)** the linkage map of chromosome 7H (Chr7H) showing positions of the ‘*raw7HS*’ locus (red bar) and flanking markers. Individual markers are shown as red dots and black horizontal bars for the whole genome and Chr7H, respectively. Genetic distances are shown in centiMorgans (cM). LOD threshold determined with a permutation test (n = 1000).

**Supplementary Fig. 5.**
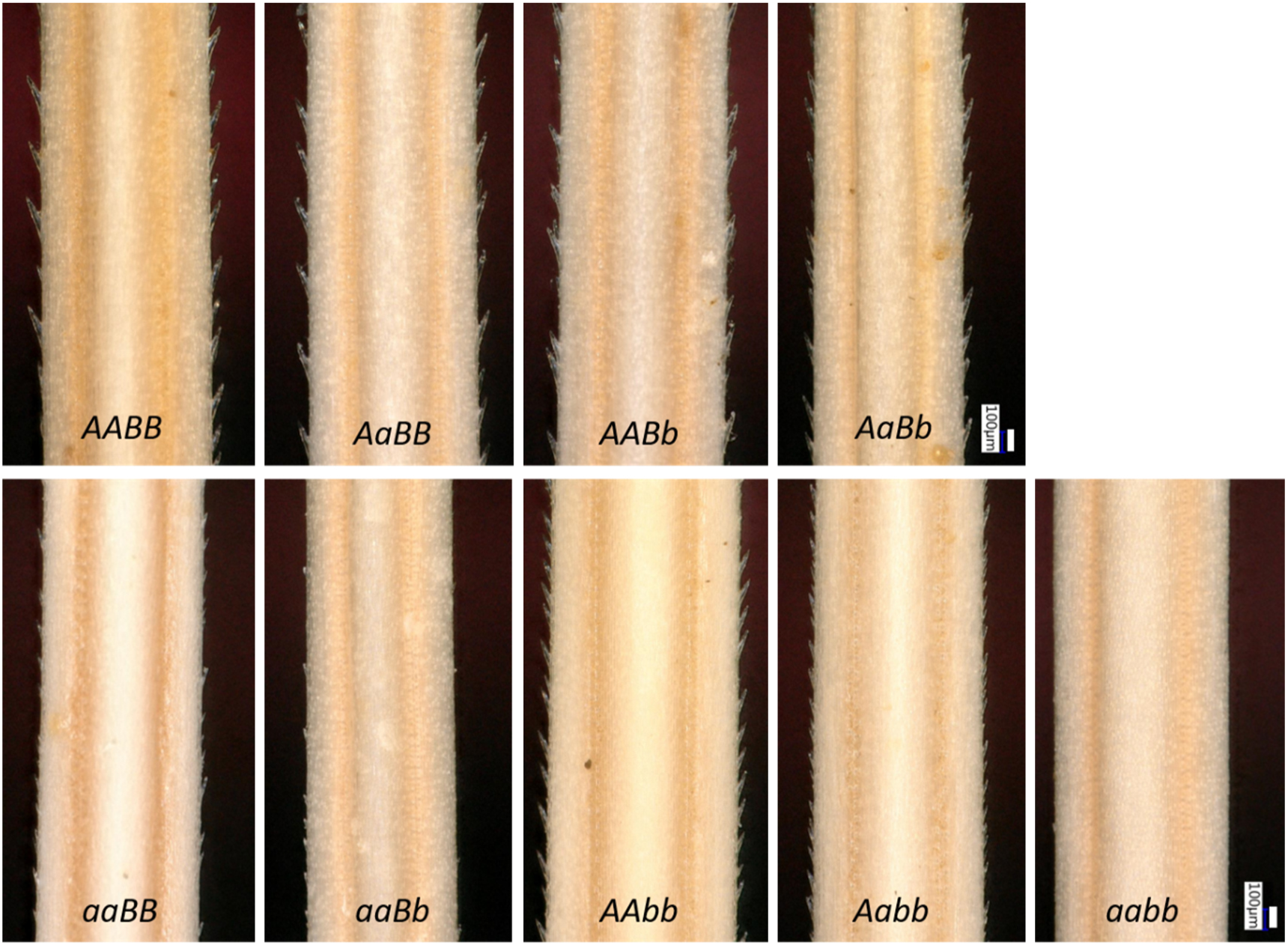
Phenotypic variation in awn roughness among all genotypic combinations of *Raw1* and *Raw7* in the Barke x MHOR 597 F_2_ population. Awn images at the basal awn position, captured using a digital microscope at 100x magnification. Genotypes reflect Wild-type (*A* = *Raw1* and *B* = *Raw7*) or mutated alleles (*a* = *raw1* and *b* = *raw7*). Scale bar = 100 μm.

**Supplementary Fig. 6.**
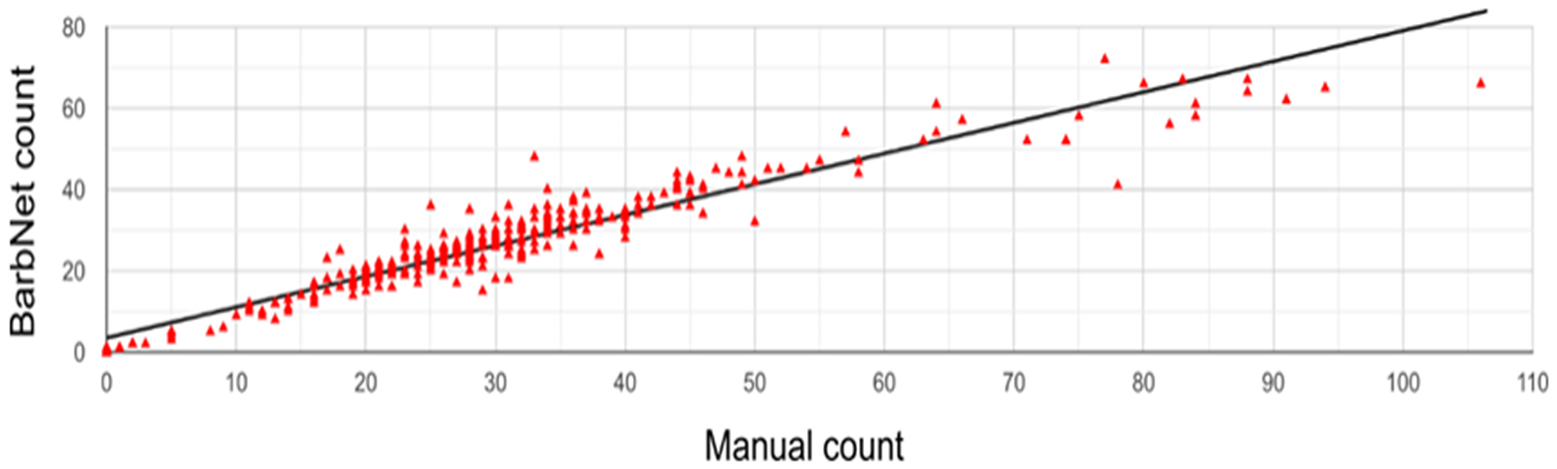
Correlation between manual and automated barb counting. Positive correlation (r = 0.96, *p* < 0.001) in barb counts quantified manually (Manual count) and via high-throughput automated counting (BarbNet count) in selected 162 F_2_ individuals of the Barke x MHOR 597 population.

**Supplementary Table 1.** Densitometric nuclear DNA content assessment via comparative absolute fluorescence intensities in barbs of cv. Morex and cv. Barke.

| Cell Type | cv. Barke (Mean ± SD) | cv. Morex (Mean ± SD) |
| --- | --- | --- |
| Stomata nuclei | 2805 ± 421 (n = 96) | 2825 ± 537 (n = 118) |
| Barb nuclei | 5239 ± 662 (n = 27) | 3460 ± 892 (n = 29) |

n = number of nuclei analyzed.

**Supplementary Table 2.**
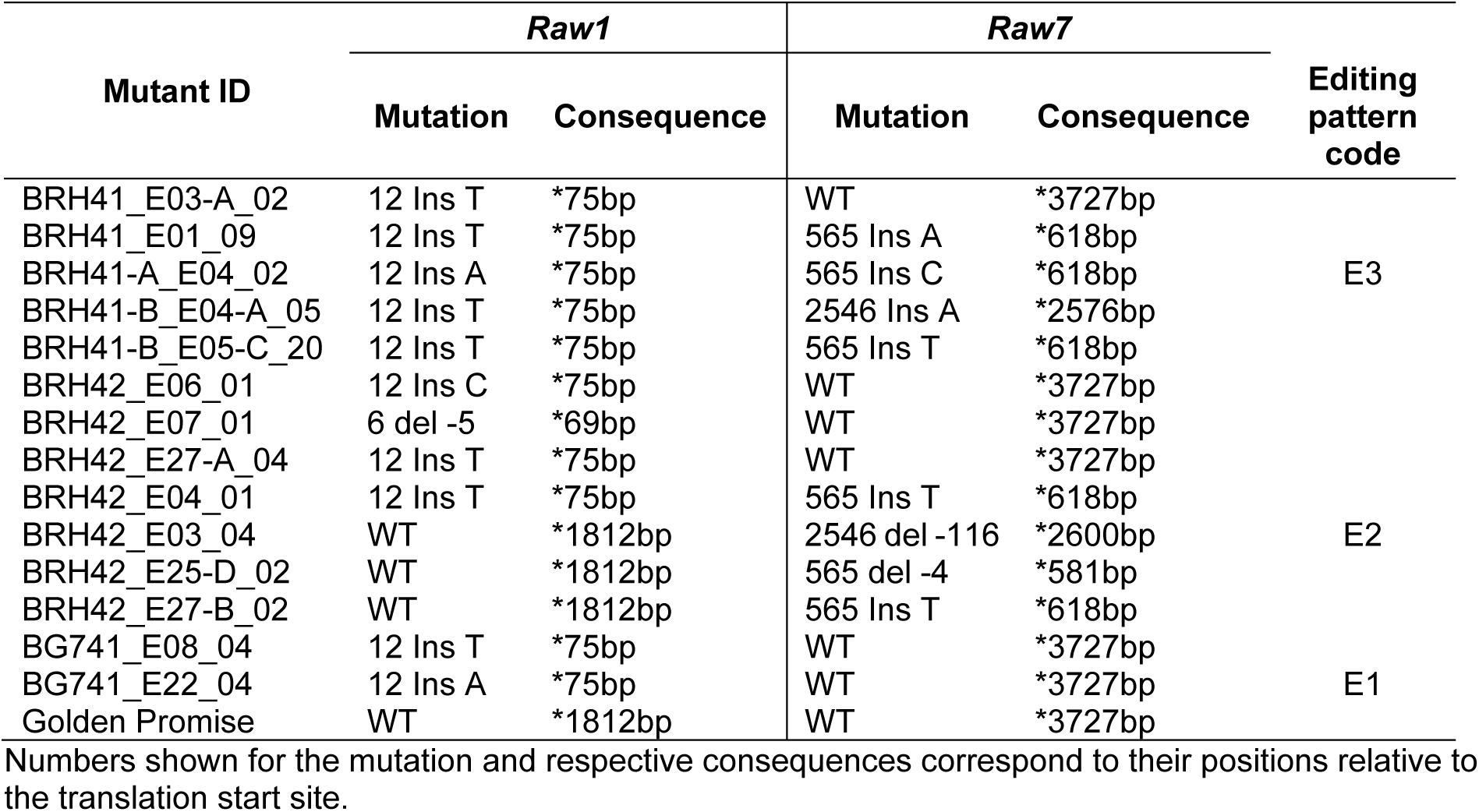
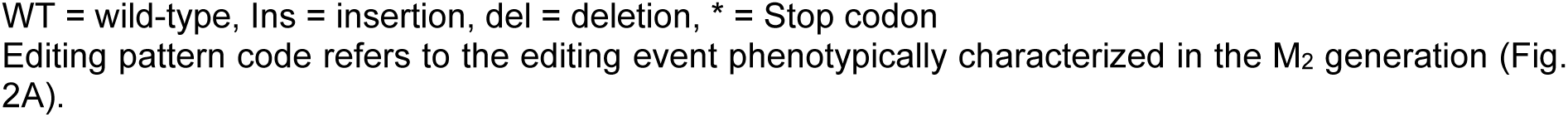
Summary of Cas9-induced homozygous single and double mutants for *Raw1* and *Raw7*.

**Supplementary Table 3.** ANOVA analyses evaluating epistatic interactions between *Raw1* and *Raw7* for their impact on barb density at basal awn position in Barke x MHOR 597 F_2_ population.

| Source of variation | df | Sum Sq | Mean Sq | F-value | Pr (>F) | Signif. |
| --- | --- | --- | --- | --- | --- | --- |
| *Ad-A* | 1 | 14791 | 14790.5 | 189.99 | < 2.2e-16 | *** |
| *Dom-A* | 1 | 4264.4 | 4264.4 | 54.78 | 8.39E-12 | *** |
| *Ad-B* | 1 | 1588.4 | 1588.4 | 20.4 | 1.25E-05 | *** |
| *Dom-B* | 1 | 2423.7 | 2423.7 | 31.13 | 1.07E-07 | *** |
| *Ad-A: Ad-B* | 1 | 7681.7 | 7681.7 | 98.67 | < 2.2e-16 | *** |
| *Ad-A: Dom-B* | 1 | 26.5 | 26.5 | 0.34 | 0.56 |  |
| *Dom-A: Ad-B* | 1 | 1712.9 | 1712.9 | 22 | 5.99E-06 | *** |
| *Dom-A: Dom-B* | 1 | 69.6 | 69.6 | 0.89 | 0.35 |  |
| Residuals | 153 | 11911 | 77.8 |  |  |  |

Signif.= Significance codes: 0 '\*\*\*' 0.001 '\*\*' 0.01 '\*' 0.05 '.' 0.1 ' ' 1, df= degree of freedom, Sq= Square, Ad = Additive, Dom = Dominance, A = *Raw1*, B = *Raw7*

**Supplementary Table 4.** ANOVA analyses evaluating epistatic interactions between *Raw1* and *Raw7* for their impact on average barb size (in pixels) at basal awn position in Barke x MHOR 597 F_2_ population.

| Source of variation | df | Sum Sq | Mean Sq | F-value | Pr (>F) | Signif. |
| --- | --- | --- | --- | --- | --- | --- |
| *Ad-A* | 1 | 4209032 | 4209032 | 283.59 | < 2.2e-16 | *** |
| *Dom-A* | 1 | 287992 | 287992 | 19.4 | 1.98E-05 | *** |
| *Ad-B* | 1 | 3397675 | 3397675 | 228.93 | < 2.2e-16 | *** |
| *Dom-B* | 1 | 39493 | 39493 | 2.66 | 0.11 |  |
| *Ad-A: Ad-B* | 1 | 90140 | 90140 | 6.07 | 0.01 | * |
| *Ad-A: Dom-B* | 1 | 68164 | 68164 | 4.59 | 0.034 | * |
| *Dom-A: Ad-B* | 1 | 122746 | 122746 | 8.27 | 0.005 | ** |
| *Dom-A: Dom-B* | 1 | 471 | 471 | 0.03 | 0.86 |  |
| Residuals | 153 | 2270783 | 14842 |  |  |  |

Signif.= Significance codes: 0 '\*\*\*' 0.001 '\*\*' 0.01 '\*' 0.05 '.' 0.1 ' ' 1, df= degree of freedom, Sq= Square, Ad = Additive, Dom = Dominance, A = *Raw1*, B = *Raw7*

**Supplementary Table 5.**
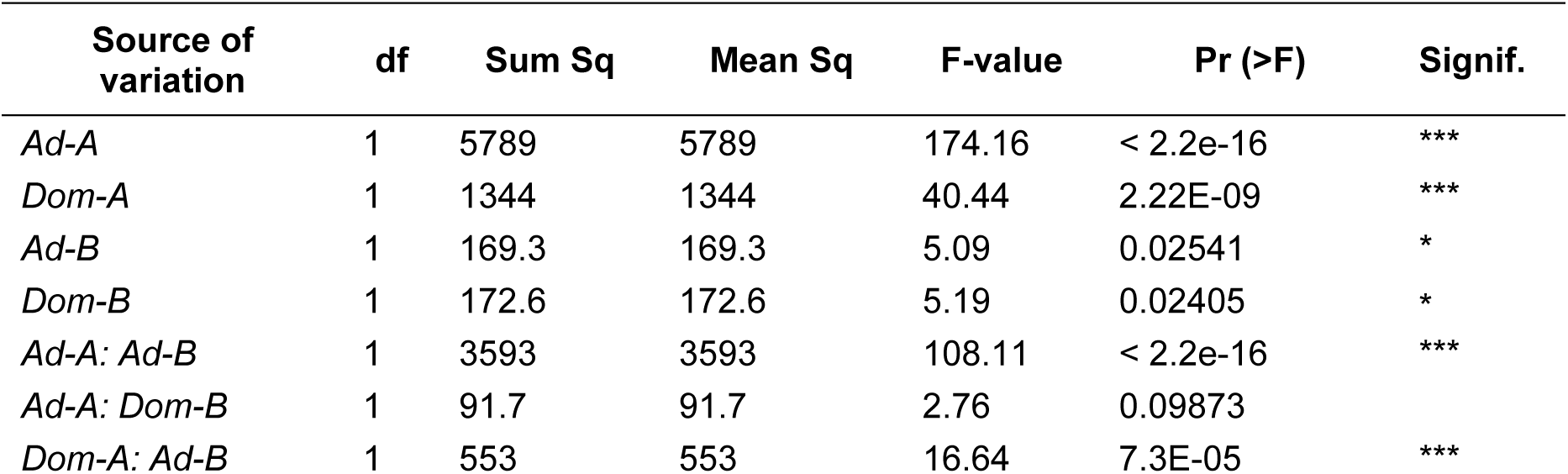
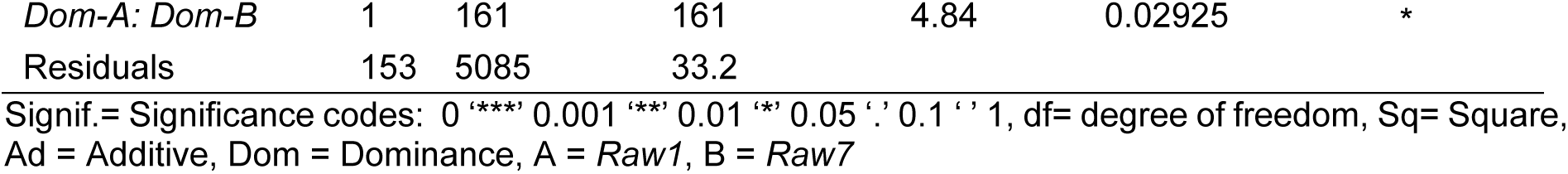
ANOVA analyses evaluating epistatic interactions between *Raw1* and *Raw7* for their impact on barb density at central awn position in the Barke x MHOR 597 F_2_ population.

**Supplementary Table 6.**
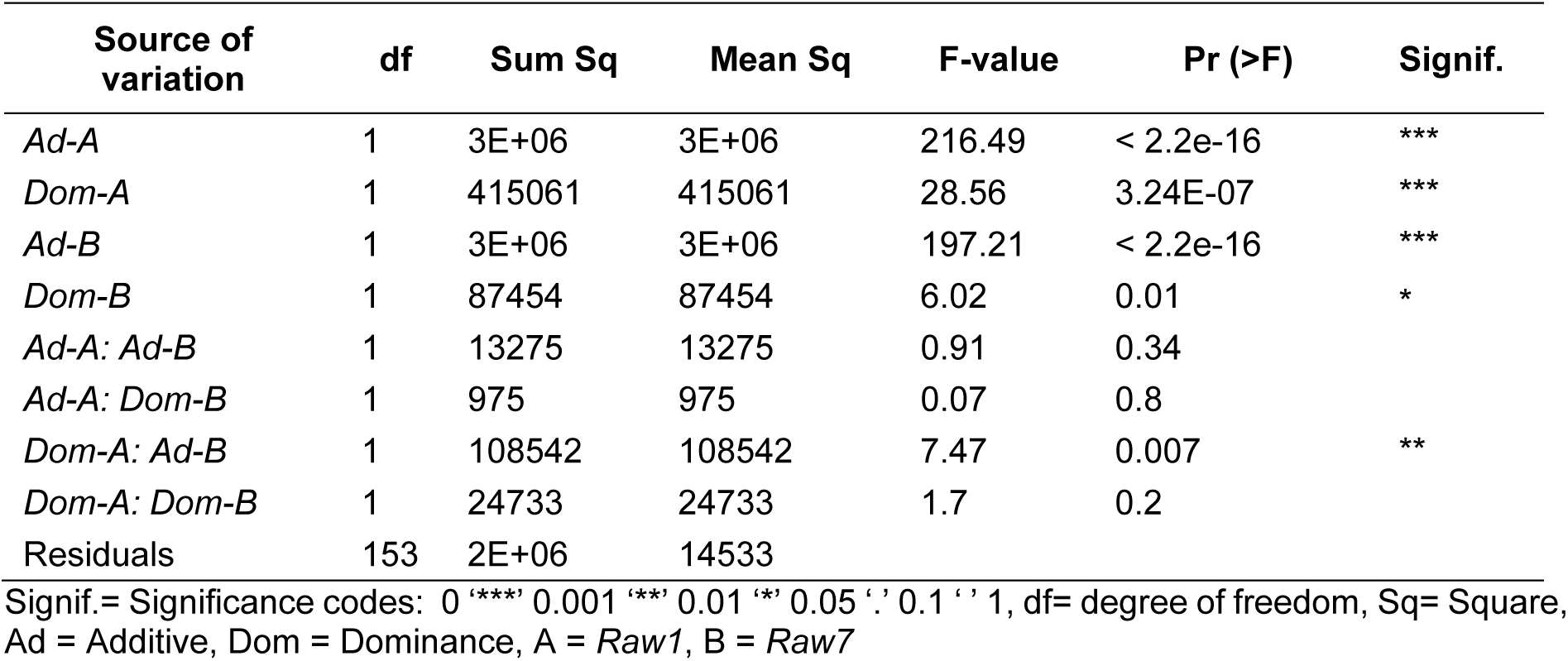
ANOVA analyses evaluating epistatic interactions between *Raw1* and *Raw7* for their impact on average barb size (in pixels) at central awn position in Barke x MHOR 597 F_2_ population.

| Source of variation | df | Sum Sq | Mean Sq | F-value | Pr (>F) | Signif. |
| --- | --- | --- | --- | --- | --- | --- |
| *Ad-A* | 1 | 3E+06 | 3E+06 | 216.49 | < 2.2e-16 | *** |
| *Dom-A* | 1 | 415061 | 415061 | 28.56 | 3.24E-07 | *** |
| *Ad-B* | 1 | 3E+06 | 3E+06 | 197.21 | < 2.2e-16 | *** |
| *Dom-B* | 1 | 87454 | 87454 | 6.02 | 0.01 | * |
| *Ad-A: Ad-B* | 1 | 13275 | 13275 | 0.91 | 0.34 |  |
| *Ad-A: Dom-B* | 1 | 975 | 975 | 0.07 | 0.8 |  |
| *Dom-A: Ad-B* | 1 | 108542 | 108542 | 7.47 | 0.007 | ** |
| *Dom-A: Dom-B* | 1 | 24733 | 24733 | 1.7 | 0.2 |  |
| Residuals | 153 | 2E+06 | 14533 |  |  |  |

Signif.= Significance codes: 0 '\*\*\*' 0.001 '\*\*' 0.01 '\*' 0.05 '.' 0.1 ' ' 1, df= degree of freedom, Sq= Square, Ad = Additive, Dom = Dominance, A = *Raw1*, B = *Raw7*

**Supplementary Table 7.**
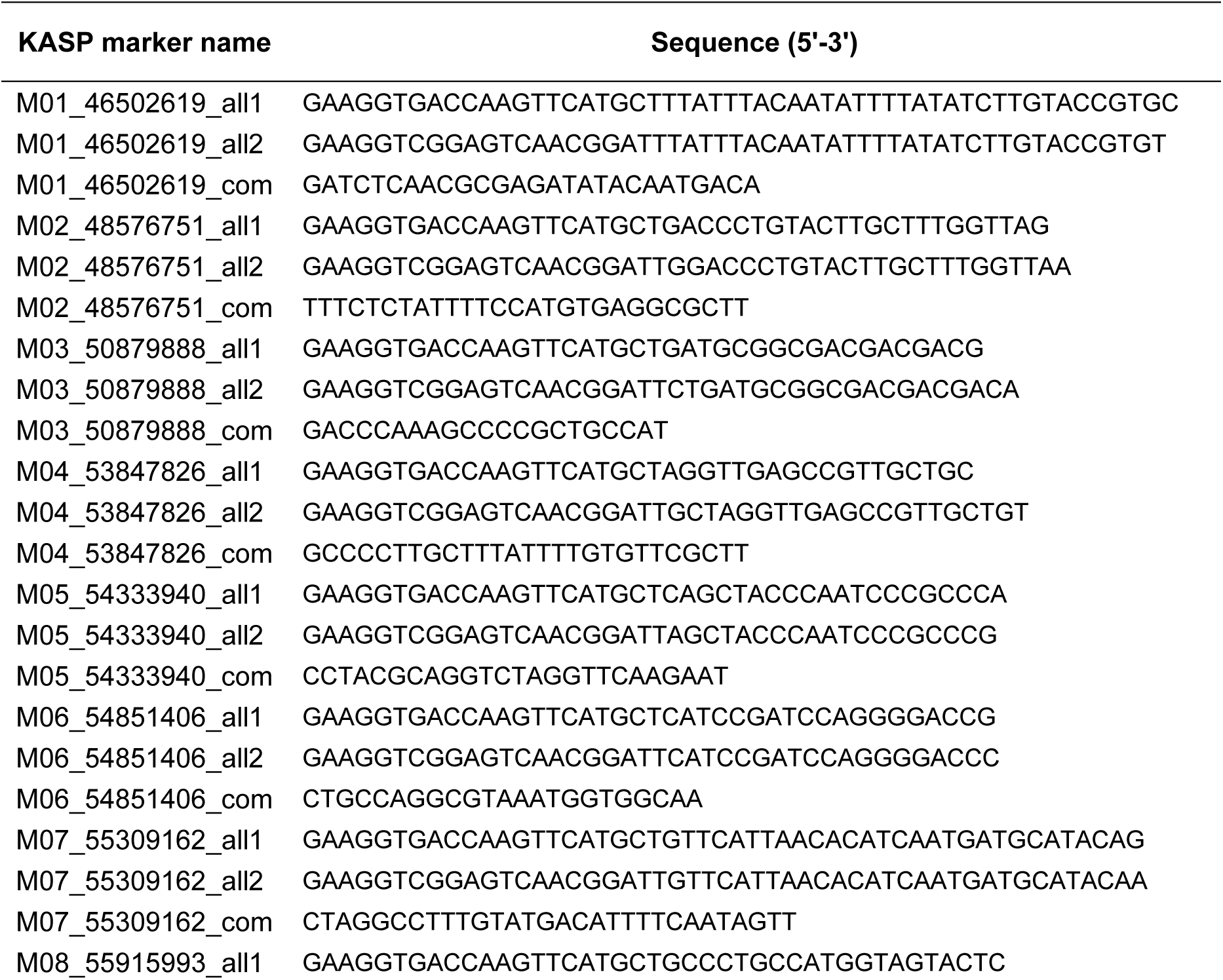
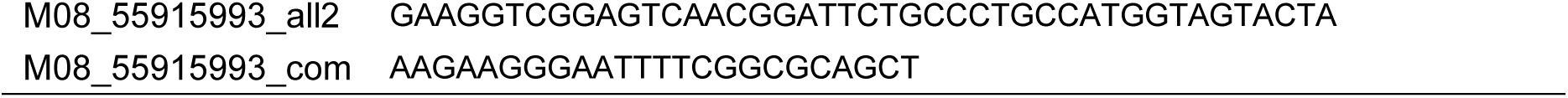
List of KASP markers and sequences used for high-resolution mapping.

**Supplementary Table 8.**
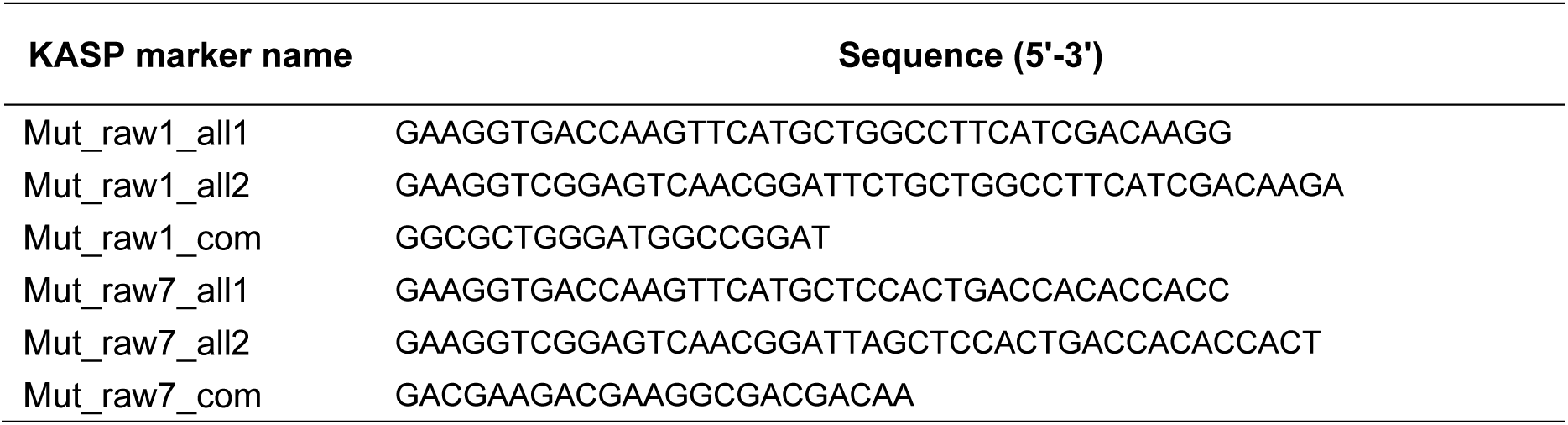
List of KASP markers and sequences used for genotyping *raw1* and *raw7* mutant alleles.

**Supplementary Table 9.**
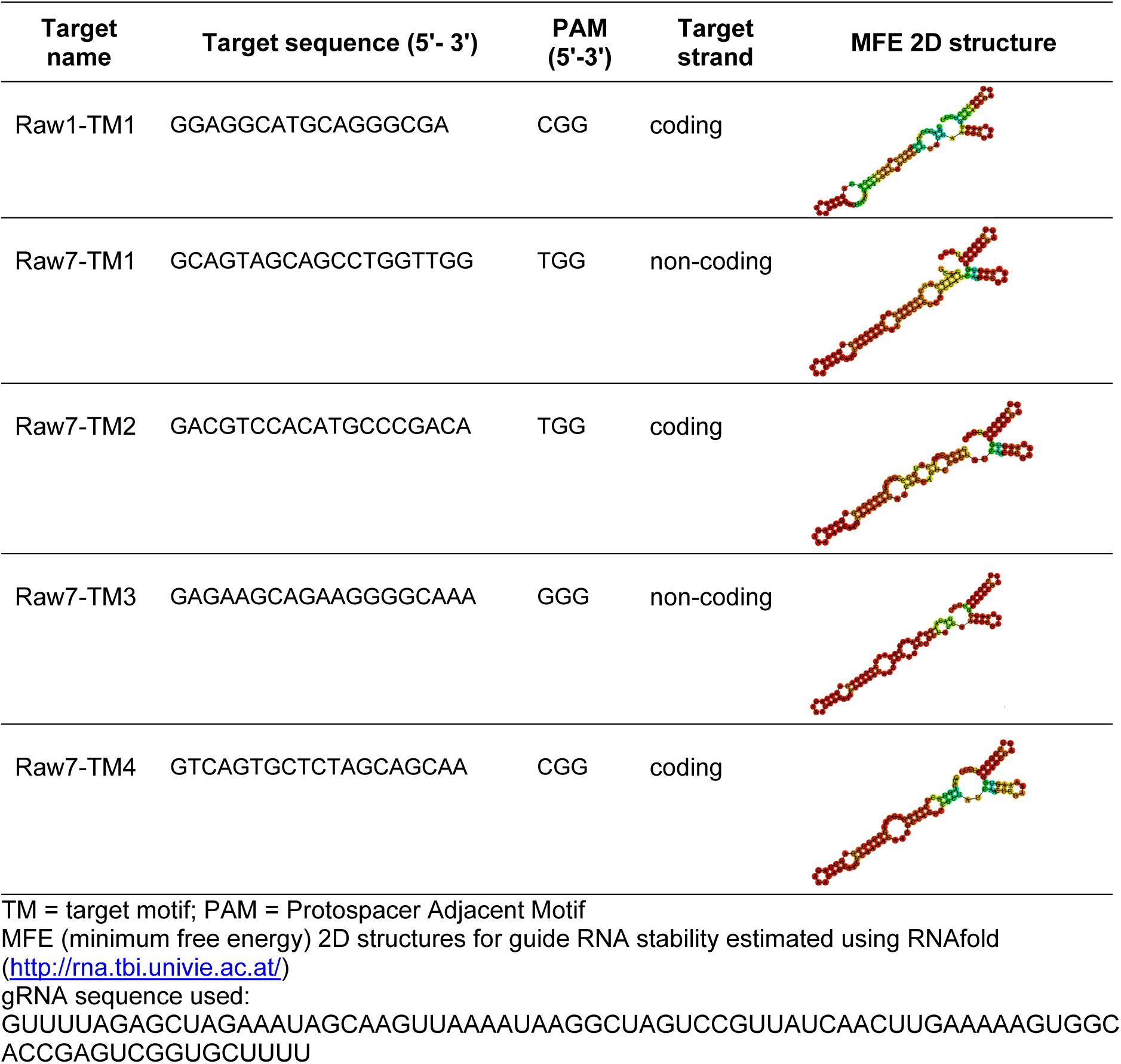
Nucleotide sequences of the target motifs and respective guide RNA (gRNA) structures selected for Cas9-mediated targeted mutagenesis of *Raw1* and *Raw7*.

**Supplementary Table 10.**
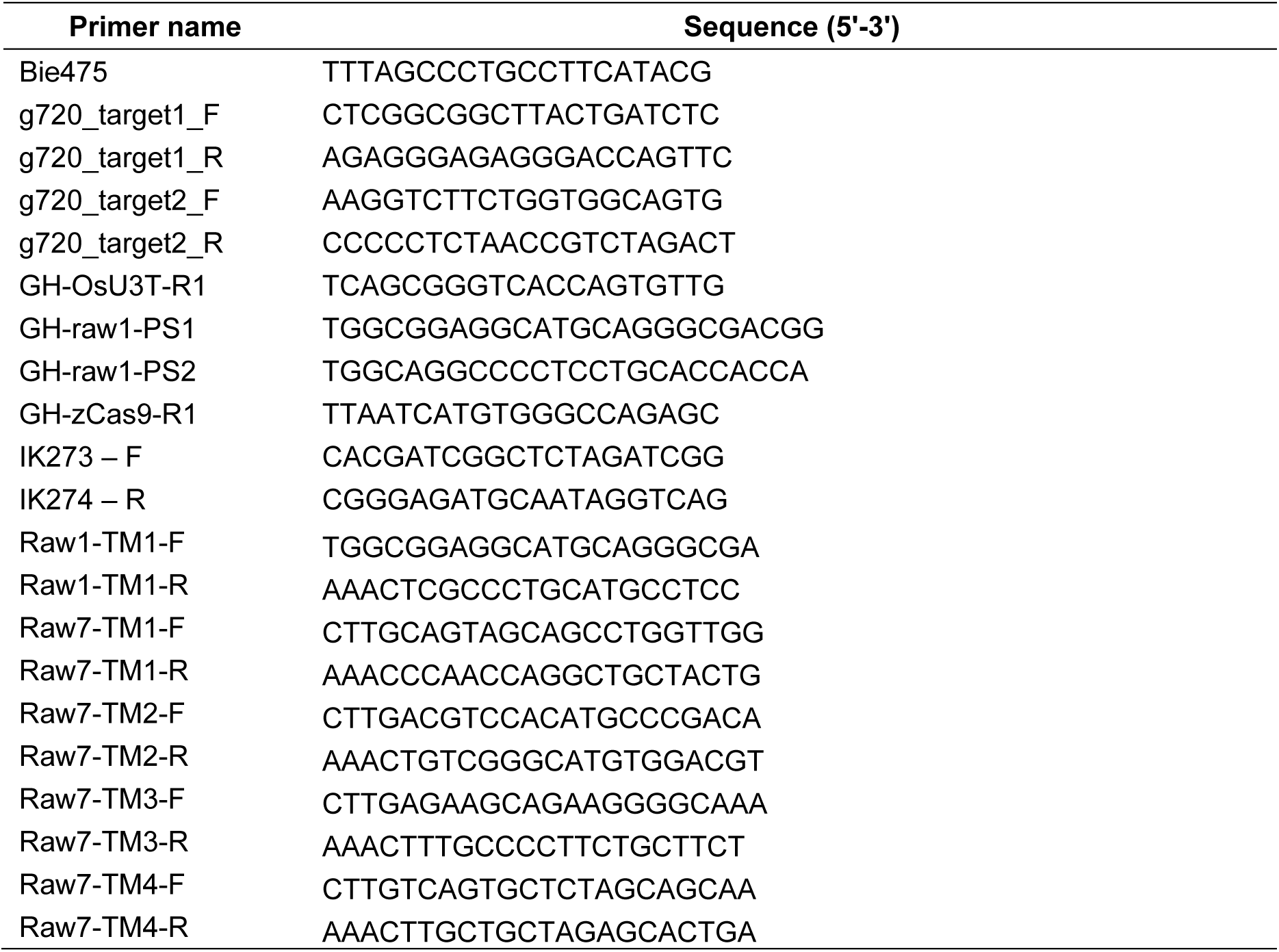
List of oligonucleotides used as primers for genotyping for editing patterns in Cas9-induced *Raw* mutants or for cloning of gRNA coding sequences.

**Supplementary Table 11.**
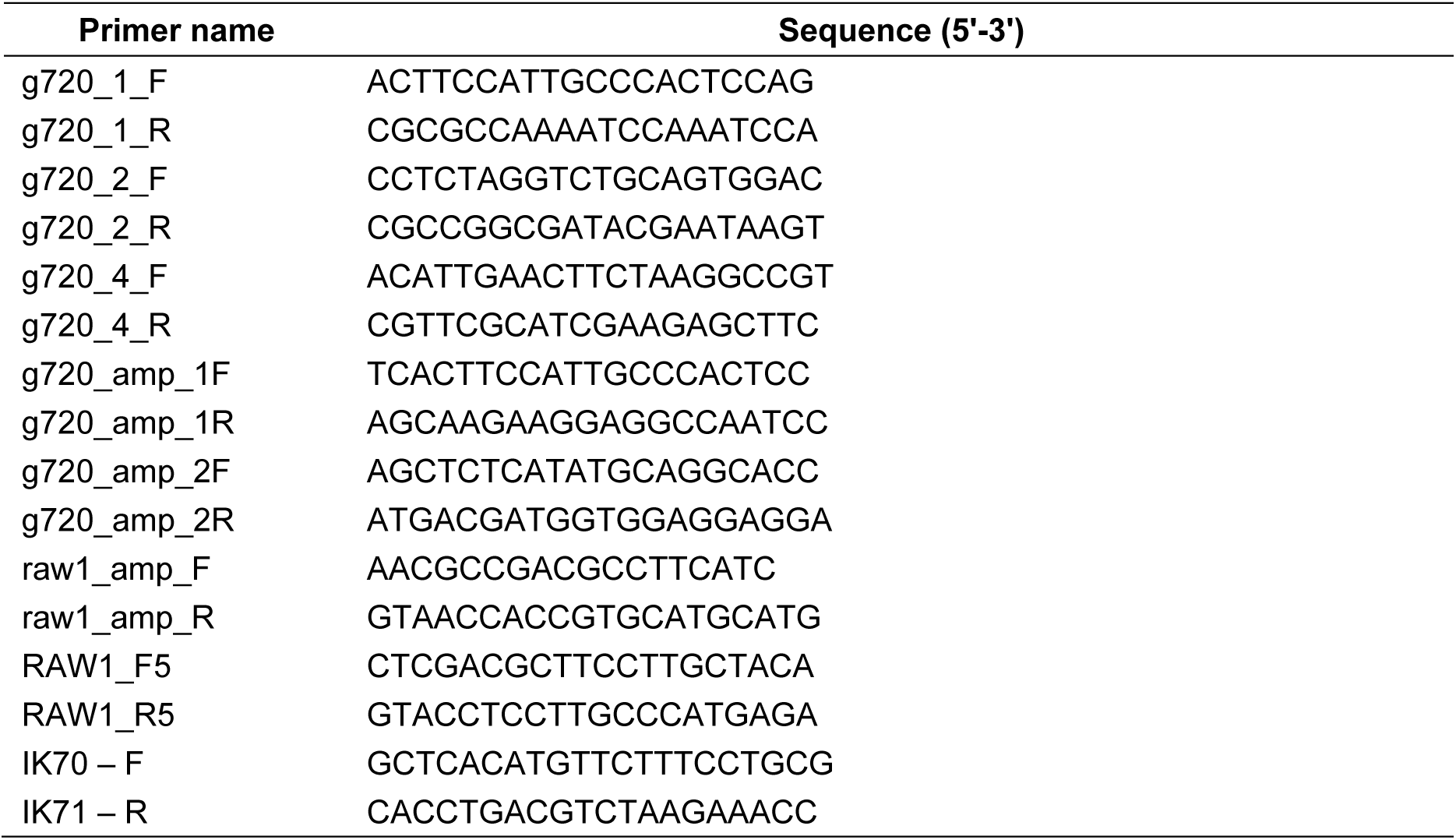
List of primers used for *Raw1* and *Raw7* CRISPR target amplification and Sanger sequencing.

